# A CLIC1 network coordinates matrix stiffness and the Warburg effect to promote tumor growth in pancreatic cancer

**DOI:** 10.1101/2023.10.27.564288

**Authors:** Jia-Hao Zheng, Yu-Heng Zhu, Jian Yang, Pei-Xuan Ji, Rui-Kang Zhao, Zong-Hao Duan, Hong-Fei Yao, Qin-Yuan Jia, Yi-Fan Yin, Li-Peng Hu, Qing Li, Shu-Heng Jiang, Yan-Miao Huo, Wei Liu, Yong-Wei Sun, De-Jun Liu

## Abstract

**BACKGROUND & AIMS:** PDAC is characterized by significant matrix stiffening and reprogrammed glucose metabolism, particularly the Warburg effect. However, it is not clear the connection between matrix stiffness and the Warburg effect and the mechanisms of action in tumor progression.

**METHODS:** The relationship between matrix stiffness and the Warburg effect was investigated from clinical, cellular, and bioinformatical perspectives. The ChIP and luciferase reporter gene assays were used to clarify the regulation mechanism of matrix stiffness on the expression of CLIC1. The expression profile and clinical significance of CLIC1 were determined in GEO datasets and a TMA. Loss-of-function and gain-of-function technics were used to determine the *in vitro* and *in vivo* functions of CLIC1. GSEA and western blotting revealed the underlying molecular mechanisms.

**RESULTS:** PDAC matrix stiffness is closely associated with the Warburg effect, and CLIC1 is a key molecule connecting tumor matrix stiffness and the Warburg effect. Increased CLIC1 expression induced by matrix stiffness correlates with poor prognosis in PDAC. CLIC1 acts as a promoter of glycolytic metabolism and facilitates tumor growth in a glycolysis-dependent manner. Mechanistically, CLIC1 inhibits the hydroxylation of HIF1α via ROS, which then increases the stability of HIF1α. Collectively, PDAC cells can sense extracellular matrix stiffness and upregulate the expression of CLIC1, which facilitates the Warburg effect through ROS/HIF1α signaling, thereby supporting tumor growth.

**CONCLUSIONS:** In the context of tumor therapy, targeted approaches can be considered from the perspectives of both extracellular matrix stiffness and tumor metabolism, of which CLIC1 is one of the targets.

## INTRODUCTION

The untransformed extracellular matrix (ECM) has the crucial supportive function of maintaining homeostasis in response to injury through its immune function and vascular and connective tissue components. However, cancer hijacks this physiological response and induces changes in the surrounding tissue matrix, creating a favorable tumor microenvironment (TME) for its rapid growth^1, 2^. The ECM, the most abundant noncellular solid component of the TME, not only maintains the three-dimensional morphological structure of tumor tissue but also influences the biological properties of tumor cells through biochemical or biophysical signals^3^. The deposition, remodeling, and cross-linking of ECM proteins in solid tumors lead to the mechanical and physical characteristics of matrix stiffening^4^. Pancreatic ductal adenocarcinoma (PDAC), as an ECM-rich cancer type, highlights this feature^5, 6^. The dense ECM not only physically obstructs the vascular system and limits the delivery of intravenous therapeutic drugs, but also induces hypoxia and suppresses tumor immunity^2, 5^. Therefore, poor patient prognosis is correlated with the highly fibrotic nature of PDAC^7^.

The role of the matrix in tumor cell invasion and metastasis has been well documented^8, 9^, but its biophysical signaling effects have been less studied. The stiffness of the ECM is one of the most prominent mechanical and physical characteristics of solid tumors. It alters the homeostatic forces on the cell surface and transmits mechanical signals generated by exogenous stroma to the cells, which in turn induces the biological behavior of cells and activation of internal signaling pathways, ultimately affecting the epithelial mesenchymal transformation, metabolic changes, invasion, and metastasis of tumor cells^10, 11^.

Cell metabolism has recently evolved as one of the processes regulated by mechanical signals^12^. Metabolic reprogramming to meet the energy and material requirements of tumors is one of the core features of tumors^13^. In this process, tumor cells tend to derive energy through glycolysis even under adequate oxygen supply conditions, which is known as the Warburg effect^14^. Moreover, the Warburg effect is more relevant to foster the survival and proliferation of PDAC cells in tumor niches with a mesenchyme-abundant TME^15^. Moreover, targeting the Warburg effect can effectively curb tumor progression^16^. Therefore, elucidating the core regulators of the Warburg effect process is essential for the development of therapeutic targets for PDAC. However, the role of the TME, especially ECM stiffness, in the Warburg effect process in PDAC has not been fully clarified.

Chloride intracellular channel 1 (CLIC1) is a member of the chloride intracellular channel (CLIC) protein family, which not only has potential ion channel activity, but also may be involved in intracellular oxidative stress, cell signaling, acidification and cell cycle regulation^17, 18^. Strikingly, accumulating studies have highlighted the role of CLIC1 in cancer. CLIC1 is highly expressed in a variety of tumor tissues^19–21^ and has been demonstrated to be an oncogenic protein that can facilitate immune escape and tumor metastasis. For instance, acetylation-stabilized CLIC1 plays a protumorigenic role in cervical cancer cells via NF-κB activation^19^. Furthermore, CLIC1 coordinates the spatial and temporal formation of lamellipodia and invadopodia through the PIP5K/PIP_2_/talin/integrin/adhesion signaling pathway to promote tumor invasion and metastasis^22^. Nevertheless, the underlying mechanisms of mediating CLIC1 within the TME remain largely unknown.

Here, we describe that CLIC1 functions as a bridge connecting tumor matrix stiffness to the Warburg effect in PDAC. We found that tumor cells can sense ECM stiffness and activate the Wnt/β-Catenin/TCF4 signaling pathway, leading to upregulation of CLIC1 expression. CLIC1 is a key regulator implicated in the Warburg effect and promotes glycolysis-dependent tumor growth. Collectively, this study reveals a previously unprecedented intersection between tissue mechanics, mechanotransduction signals, and metabolic remodeling in PDAC.

## MATERIALS & METHODS

### Clinical samples

Three cohorts from Ren Ji Hospital were included in the study: (I) the tumor micro array (TMA) cohort (n = 107), (II) the proteomics cohort with positron emission tomography-computed tomography (PET-CT) information (n = 8), and (III) the stiffness-glycolysis cohort (n = 14). The clinical samples were obtained from patients with PDAC treated surgically and confirmed by postoperative pathology at Ren Ji Hospital, and each patient provided written informed consent. Before surgery, none of the patients had received radiotherapy or chemotherapy. Pathological information and PET-CT data were collected from the medical record system of Ren Ji Hospital for all enrolled patients. The follow-up time was counted from the date of surgery to the date of PDAC-related death or the last known date of follow-up. This study was approved by the Research Ethics Committee of Ren Ji Hospital, School of Medicine, Shanghai Jiao Tong University (approval number: RA-2019-116).

### Statistical analysis

Data management and analysis were carried out using GraphPad Prism 8 and SPSS 22.0. Continuous variables are presented as the mean ± standard error of the mean (SEM) and were compared using independent Student’s t test or one-way ANOVA. Correlations between continuous variables were assessed using Pearson’s correlation test or Spearman’s rho test, while categorical variables were assessed using the chi-square test. For survival analyses, we used the KaplanLMeier method to compare the correlation between variables and overall survival, and the log-rank test to analyze survival curves. The Cox regression model was used to conduct univariable and multivariable analyses. For the results, P < 0.05 was considered statistically significant.

## RESULTS

### Relationship between ECM stiffness and glucose metabolism in PDAC

To explore our conjecture that extracellular matrix components or properties are linked to glucose metabolism, we first investigated this at the bioinformatics level. Starting from the glycolysis-related gene sets, we downloaded four commonly used gene signatures from the MsigDB database. A cancer-associated ECM (C-ECM) gene signature^23^ was used to define the tumor matrix-related gene set and analyze its correlation with glycolysis. It was found that there was a significant positive correlation between them (Figure 1A). To further elucidate the association between matrix stiffness and glucose metabolism, we used picrosirius red (PSR) staining and immunohistochemical (IHC) staining of collagen type I to evaluate the stiffness level of PDAC tissues from a clinical perspective. According to the highly specific characteristics of PSR staining birefringence on collagen fibers under a polarizing microscope, the degree of fibrillar collagen in tissue sections was observed and analyzed, combined with the immunohistochemical staining of collagen type I. Based on the threshold of fibrillar collagen, the patients were divided into a high-stiffness group (n = 7) and a low-stiffness group (n = 7). IHC analysis showed that three key glycolytic components (HK2, GLUT1 and LDHA) had higher expression in PDAC samples with high stiffness than in those with low stiffness (Figure 1B-C). Moreover, based on the standardized uptake value-max (SUV-max) of PET-CT reflecting the glucose uptake ability in the local PDAC tumor niche, we also compared the metabolic status of glucose in PDAC tissues between the high- and low-stiffness groups (Figure 1D). SUV-max tended to be higher in the high-stiffness group than in the low-stiffness group (Figure 1E). Subsequently, we intended to validate our findings at the cellular level. We used polyacrylamide gels to simulate two different stiffnesses of the extracellular matrix (0.5 kPa and 12 kPa) to culture commonly used PDAC cells *in vitro* and examined the differences in mRNA expression of three genes (*HK2, GLUT1 and LDHA*) in the two groups of cells. The results were consistent, as expected, and tumor cells cultured at high stiffness levels had higher expression of the three key glycolytic components (Figure 1F).

**Figure 1.**
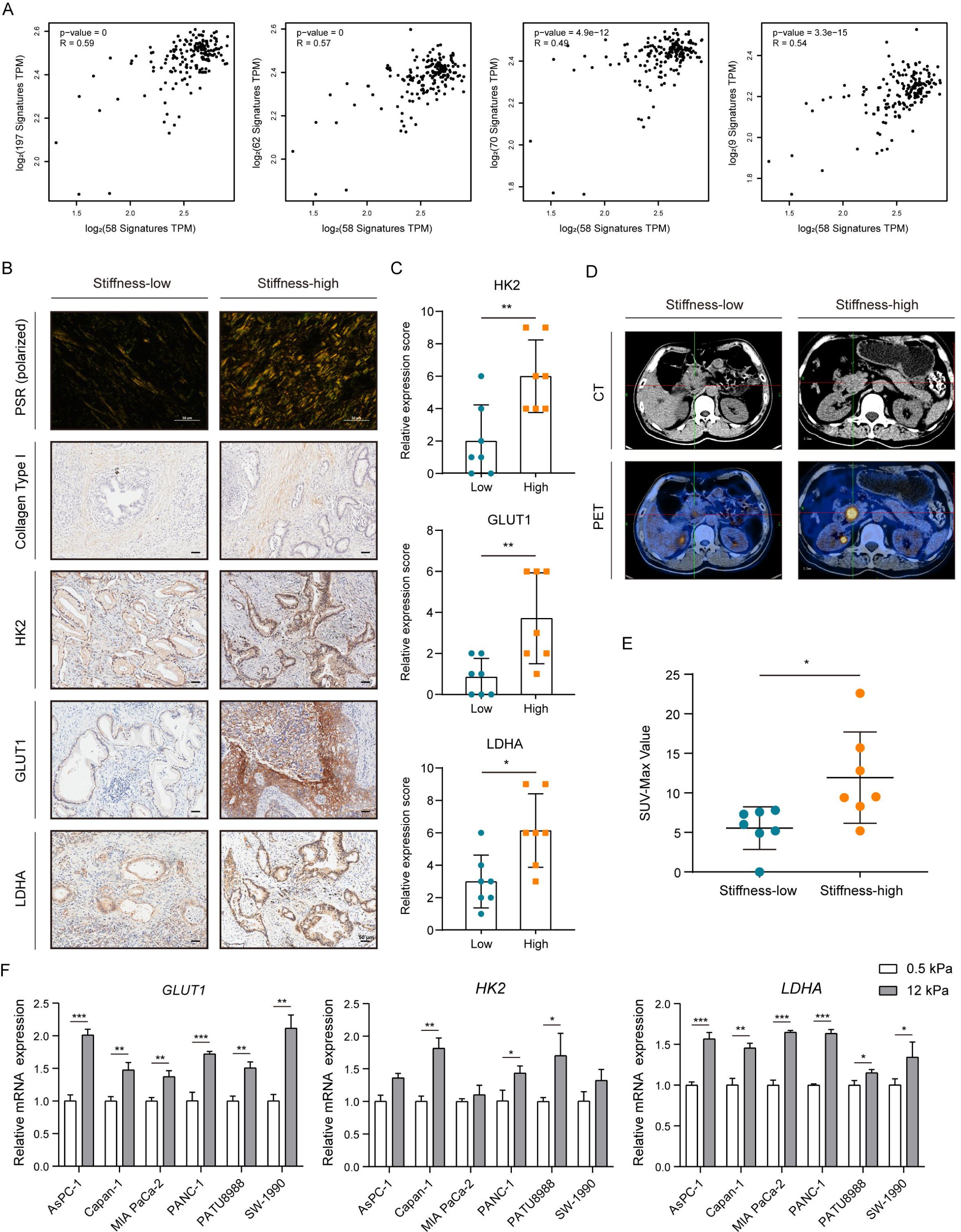
Relationship between ECM stiffness and glucose metabolism in PDAC. (A) Correlation analysis between the C-ECM gene set and four commonly used glycolysis-related gene sets (HALLMARK_GLYCOLYSIS, KEGG_GLYCOLYSIS_GLUCONEOGENESIS, REACTOME_GLYCOLYSIS, and BIOCARTA_GLYCOLYSIS_PATHWAY). (B-C) Representative IHC maps and quantitative results of the expressions of three key glycolytic components (HK2, LDHA, and GLUT1) in PDAC samples with low and high stiffness (n = 7 cases per group, mean ± SEM., two-tailed unpaired *t* test, scale bar: 50 μm). (D-E) Comparison of PET-CT images and corresponding SUV-Max between high- and low-stiffness PDAC samples (n = 7 cases per group, mean ± SEM., two-tailed unpaired *t* test). (F) Difference in the mRNA expression of *HK2, LDHA* and *GLUT1* in soft (0.5 kPa) and stiff (12 kPa) gels (mean ± SEM., two-tailed unpaired *t* test). \**P* < 0.05, \*\**P* < 0.01, \*\*\**P* < 0.001.

### Screening and verification of tumor matrix- and glucose metabolism-correlated genes

To further explore the relationship between matrix stiffness and glycolysis, we screened for genes related to the tumor matrix and glucose metabolism. First, based on the C-ECM gene set mentioned above, we utilized gene set variation analysis (GSVA) to analyze the enrichment of the gene set in each sample in the TCGA-PAAD database and obtained GSVA scores calculated from gene expression levels enriched in the gene set. Using the median of GSVA scores as a cutoff, we divided TCGA-PAAD patients into high-score and low-score groups, and used the “limma” R package to identify differentially expressed genes (DEGs) between the two groups, as shown in Figure 2A, to identify genes associated with the tumor matrix. Next, the proteomics cohort with PET-CT information (n = 8) were divided into two groups according to the SUV-max, and gene expression in these two groups of PDAC tissues was screened for glucose metabolism-correlated DEGs with isobaric tags for relative and absolute quantitation proteomics technology using FC > 1.5 and p < 0.05 conditions (Figure 2B). Furthermore, we annotated the cellular functions of the tumor matrix-correlated genes and the glucose metabolism-correlated genes using the KEGG database (Supplementary Table 4, Supplementary Table 5). The KEGG analysis of the tumor matrix-correlated genes showed fascinating results, including “ECM-receptor interaction”, “cell adhesion molecules”, “cytokine-cytokine receptor interaction”, “focal adhesion” and “regulation of actin cytoskeleton” (Supplementary Figure 1A). And the pathway annotation of DEGs derived from proteomics is mainly focused on the metabolic direction, such as “glycine, serine and threonine metabolism”, “carbon metabolism” and “glycolysis/gluconeogenesis” (Supplementary Figure 1B). As shown in Figure 2C, we integrated four gene sets, which were obtained from (1) the DEGs identified in Figure 2A, (2) the DEGs identified in Figure 2B, (3) genes differentially expressed between tumor and normal tissues in the TCGA-PAAD database, and (4) the top 500 prognostic genes in the TCGA-PAAD database from GEPIA platform. As a result, CLIC1 was obtained through this integration process.

**Figure 2.**
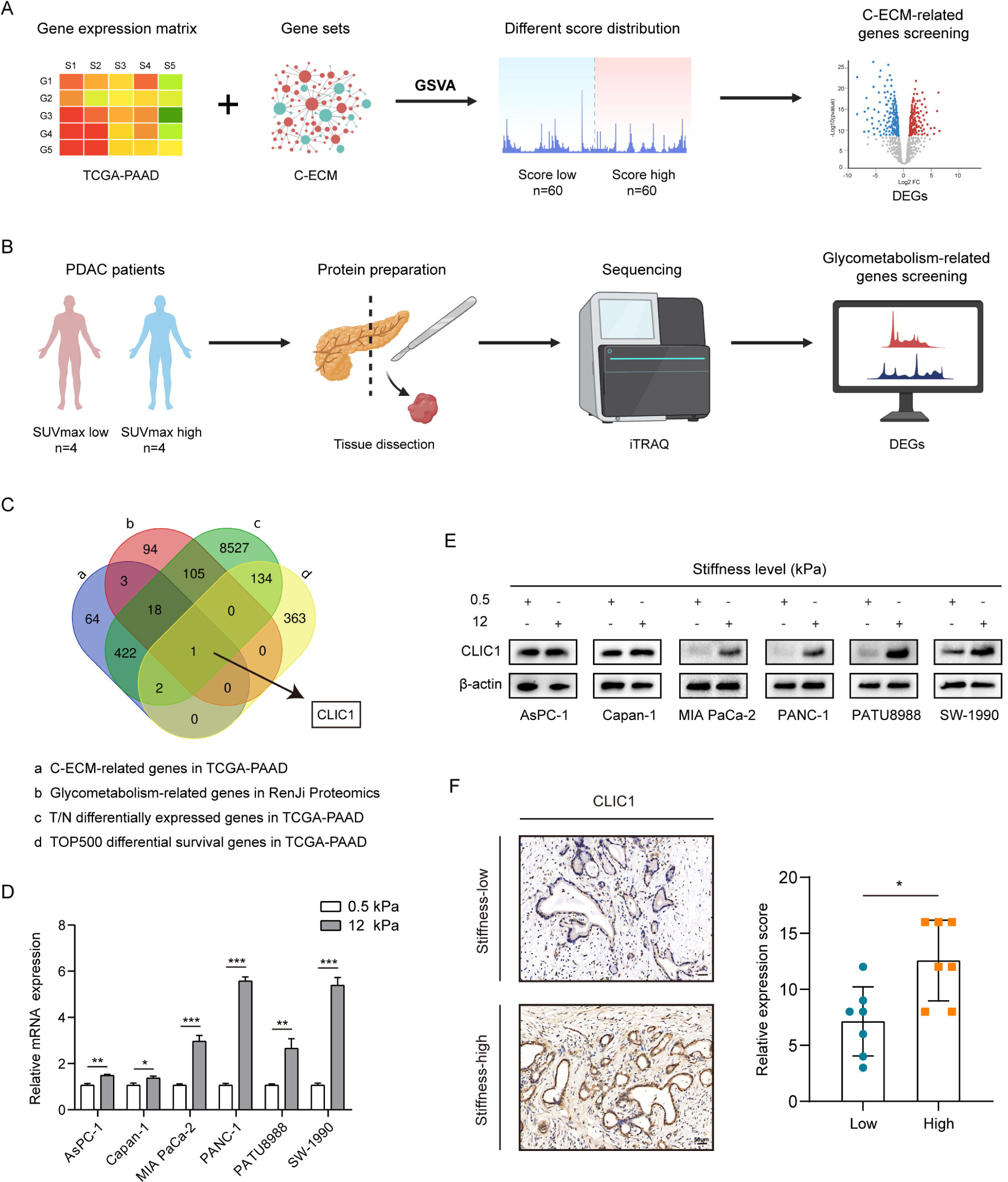
Screening and verification of tumor matrix- and glucose metabolism-correlated genes. (A) Flow chart of screening tumor matrix-related DEGs based on the TCGA-PAAD database. (B) Flow chart of screening DEGs related to glucose metabolism based on Ren Ji sample proteomics data. (C) Venn diagram for screening tumor matrix- and glycolytic-related genes. (D) Different extracellular matrix stiffnesses were simulated to explore their effects on the expression of *CLIC1* at the mRNA level (mean ± SEM., two-tailed unpaired *t* test). (E) Differential expression of CLIC1 protein levels in PDAC cells under different matrix stiffnesses. (F) Representative IHC plots and quantitative analysis of CLIC1 expression in PDAC samples from the high- and low-stiffness groups (n = 7 cases per group, mean ± SEM., two-tailed unpaired *t* test, scale bar: 50 μm). \**P* < 0.05, \*\**P* < 0.01, \*\*\**P* < 0.001.

To investigate whether ECM stiffness induces the expression of CLIC1, we also detected CLIC1 expression under different matrix stiffness conditions at both the cellular and clinical sample levels. Tumor cells were cultured with two hardness levels of gels to simulate matrix stiffness *in vitro*. Compared with the soft-gel group, as shown in Figure 2D-E, the expression of CLIC1 was induced to increase in MIA PaCa-2, PANC-1, PATU8988, and SW-1990 cells cultured in the stiff gel. Additionally, we evaluated the expression of CLIC1 by immunohistochemistry in tumor tissue with different levels of stiffness, which showed that CLIC1 expression tended to be higher in patients in the high-stiffness group (Figure 2F). The above results suggest that the expression of CLIC1 was indeed increased by the induction of matrix stiffness.

### The nuclear translocation of TCF4 induced by matrix stiffness transcriptionally regulates CLIC1 expression

In mechanotransduction, transcription factors are the most critical molecules that ultimately transmit signals triggered by ECM stiffness into the nucleus to regulate gene expression^24^. For instance, YAP/TAZ is the most well-studied transcription factor regulated by ECM stiffness^25^. To further clarify why CLIC1 expression is upregulated under high stiffness conditions, we focused on transcription factors. Two criteria were set for screening upstream transcription factors that may be affected by matrix stiffness and bind to the CLIC1 promoter region to mediate its expression: 1) transcription factors of CLIC1 predicted by the AnimalTFDB database, and 2) genes differentially expressed in the TCGA-PAAD database based on the C-ECM gene set. As shown in Figure 3A, four transcription factors, EGR2, FOXA2, SPI1, and TCF4, were obtained for subsequent studies. We then explored whether the nuclear translocation of these four transcription factors was altered in PDAC cells cultured at different stiffnesses. Immunoblot analysis of separate nuclear/cytosolic fractions indicated that nuclear localization of FOXA2 and TCF4 was increased at higher hardness levels (Figure 3B). We then found that the expression of *CLIC1* was significantly reduced after knocking down TCF4 in PANC-1 and PATU8988 cells, while interference with FOXA2 did not show a significant difference (Figure 3C). Furthermore, we conducted luciferase assays to investigate whether TCF4 directly affects CLIC1 expression. As shown in Figure 3D, the wild-type CLIC1 promoter, but not the mutant construct, was activated by TCF4 overexpression in PATU8988 cells cultured at high stiffnesses. The ChIP-PCR assay also confirmed that CLIC1 is a target gene of TCF4 (Figure 3E). TCF4 belongs to the TCF family, which is involved in the activation of the classical Wnt pathway by binding β-catenin proteins^26^. Thus, we employed a TOP/FOP flash assay to detect the activation of the Wnt/β-catenin pathway in PATU8988 cells under different substrate stiffness conditions. As shown in Figure 3F, the relative fluorescence intensity was significantly enhanced at higher stiffness levels, while the TOP/FOP reporter activity subsequently declined only with the addition of ICG-001 (an inhibitor of β-catenin/TCF-mediated transcription) but not PF-573228 (an ATP-competitive FAK inhibitor) and RGDS (an inhibitor of integrin receptor function). The results suggest that the Wnt/β-catenin/TCF4 signaling pathway mediates increased stiffness on the upregulation of CLIC1 expression.

**Figure 3.**
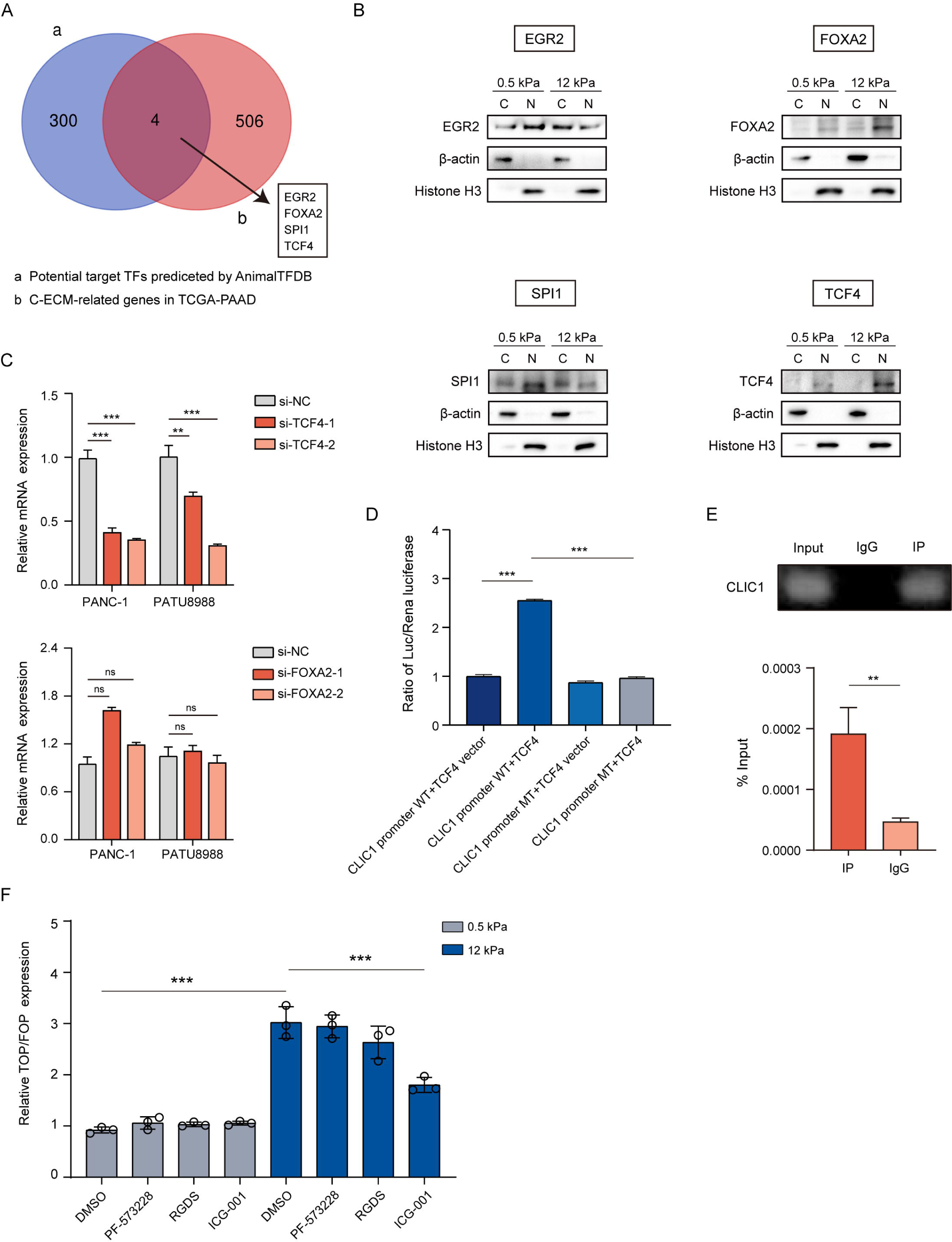
The nuclear translocation of TCF4 induced by matrix stiffness transcriptionally regulates CLIC1 expression. (A) Venn diagram screening of transcription factors that may be affected by matrix components and bind to the CLIC1 promoter region. (B) The expression of EGR2, FOXA2, SPI1 and TCF4 transcription factors in cells cultured with different matrix stiffnesses *in vitro*. (C) The difference in *CLIC1* expression after knocking down FOXA2 or TCF4 in PANC-1 and PATU8988 cell lines in the 12 kPa gel (mean ± SEM., two-tailed unpaired *t* test). (D) The luciferase activity of wild-type (WT) and mutant (MT) CLIC1 promoter reporter gene plasmids and TCF4 overexpression and control plasmids in PATU8988 cells in the 12 kPa gel (mean ± SEM., one-way ANOVA with Tukey’s multiple comparison test). (E) ChIP-PCR analysis of TCF4 binding to the CLIC1 promoter region in PATU8988 cells at 12 kPa (mean ± SEM., two-tailed unpaired *t* test). (F) Determination of the relative fluorescence intensity of TOP/FOP flashes in PATU8988 cells after treatment with DMSO, PF-573228, RGDS and ICG-001 at different levels of stiffness (mean ± SEM., one-way ANOVA with Tukey’s multiple comparison test). ns, No significance, \*\**P* < 0.01, \*\*\**P* < 0.001.

### Expression pattern and clinical relevance of CLIC1 in PDAC tissues

To gain further insight into the expression pattern of CLIC1 in PDAC, we first analyzed *CLIC1* expression in seven pancreatic cancer GEO datasets (Figure 4A), and the results showed that *CLIC1* expression was higher in tumor tissues than in adjacent normal pancreatic tissues at the mRNA level. We then verified our results at the protein level by performing IHC staining in a TMA containing 107 PDAC specimens and matched adjacent pancreatic tissue from the Ren Ji cohort. As shown in Figure 4B-C, the expression of CLIC1 was scored and counted according to staining intensity and area, and the tumor tissue showed a higher staining score. Subsequently, based on the staining scores of the TMA, we stratified patients from the TMA cohort into high- and low-CLIC1 expression groups. By integrating these data with clinical data, we discovered that CLIC1 expression was positively correlated with the TNM stage of PDAC (Figure 4D). Univariate and multivariate Cox regression analyses revealed that CLIC1 expression was an independent prognostic risk factor for PDAC (Supplementary Figure 2A, Figure 4E). Moreover, as shown in Figure 4F-I, KaplanLMeier survival analysis demonstrated a significantly prolonged median survival time in the low-expression group compared to the high-expression group, which was consistent with the results obtained from the analysis of GEO databases. We also investigated the expression pattern of CLIC1 in *Kras*^G12D^/*Trp53*^R172H^/*Pdx1*-Cre (KPC) mice. As expected, the expression of CLIC1 was elevated in pancreatic intraepithelial neoplasms (PanINs) and tumor tissues from KPC mice (Figure 4J, Supplementary Figure 2B). Taken together, these data suggest that CLIC1 is generally upregulated and may serve as a predictive indicator for malignant progression in PDAC.

**Figure 4.**
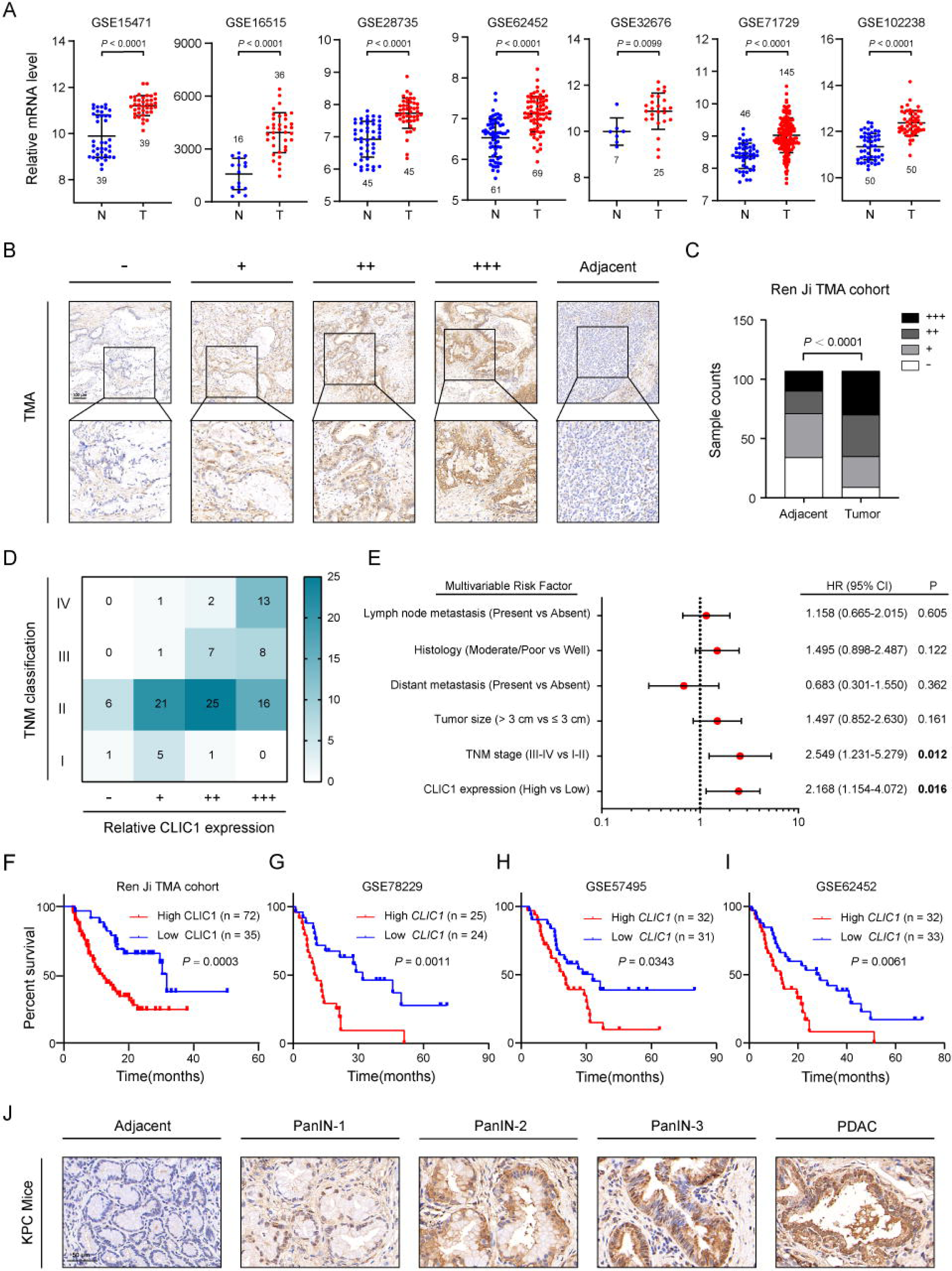
Expression pattern and clinical relevance of CLIC1 in PDAC tissues. (A) The expression of CLIC1 in tumor tissues and paired nontumor tissues in 7 GEO datasets (GSE15471, GSE16515, GSE28735, GSE62452, GSE32676, GSE71729, and GSE102238) (mean ± SEM., two-tailed unpaired *t* test). (B) Representative immunohistochemical staining images of CLIC1 expression in a TMA (n = 107 samples, 3 fields assessed per sample, scale bar: 100 μm). (C) Statistics on the staining of CLIC1 expression in cancer and adjacent tissues in the TMA (Pearson chi-squared test). (D) Heatmap of correlation analysis between staining profiles of CLIC1 expression and TNM classification according to the IHC. (E) Multivariate Cox regression analysis of clinicopathological parameters for overall survival based on the TMA cohort. Low CLIC1 expression was defined as a score of “-” or “+” and high CLIC1 expression with a score of “++” or “+++”. (F-I) Kaplan–Meier analysis of the overall survival rate of PDAC patients with different CLIC1 expression levels based on the GEO database (GSE78229, GSE57495, and GSE62452) and the Ren Ji cohort (two-sided log-rank test). CI Confidence interval, HR Hazard ratio. (J) Standard images of CLIC1 immunohistochemical staining of KPC mice at different stages of PDAC progression (n = 10 samples, 5 fields assessed per sample, scale bar: 50 μm).

### CLIC1 promotes the proliferative capacity of PDAC cells *in vivo* and *in vitro*

To probe the role of CLIC1 in the development of PDAC, we initially detected the expression of CLIC1 in 9 pancreatic cancer cell lines (Supplementary Figure 3A-B). PANC-1 and PATU8988 cells were selected for RNA interference, as they exhibited relatively high levels of CLIC1 expression (Supplementary Figure 3C-D). In addition, Capan-1 and SW-1990 cells with relatively low CLIC1 expression were selected for exogenous overexpression (Supplementary Figure 3E-F). We then assessed the impact of CLIC1 on cell growth using a CCK-8 assay. Our results revealed a significant reduction in the proliferative capacity of PANC-1 and PATU8988 cells upon downregulation of CLIC1 expression, while overexpression of CLIC1 in Capan-1 and SW-1990 cells led to a significant increase in their proliferation (Figure 5A-B). These findings were further corroborated by colony formation assays, which demonstrated a similar trend (Figure 5C-D). Therefore, our data strongly suggest that CLIC1 plays a crucial role in promoting the proliferative potential of PDAC cells.

**Figure 5.**
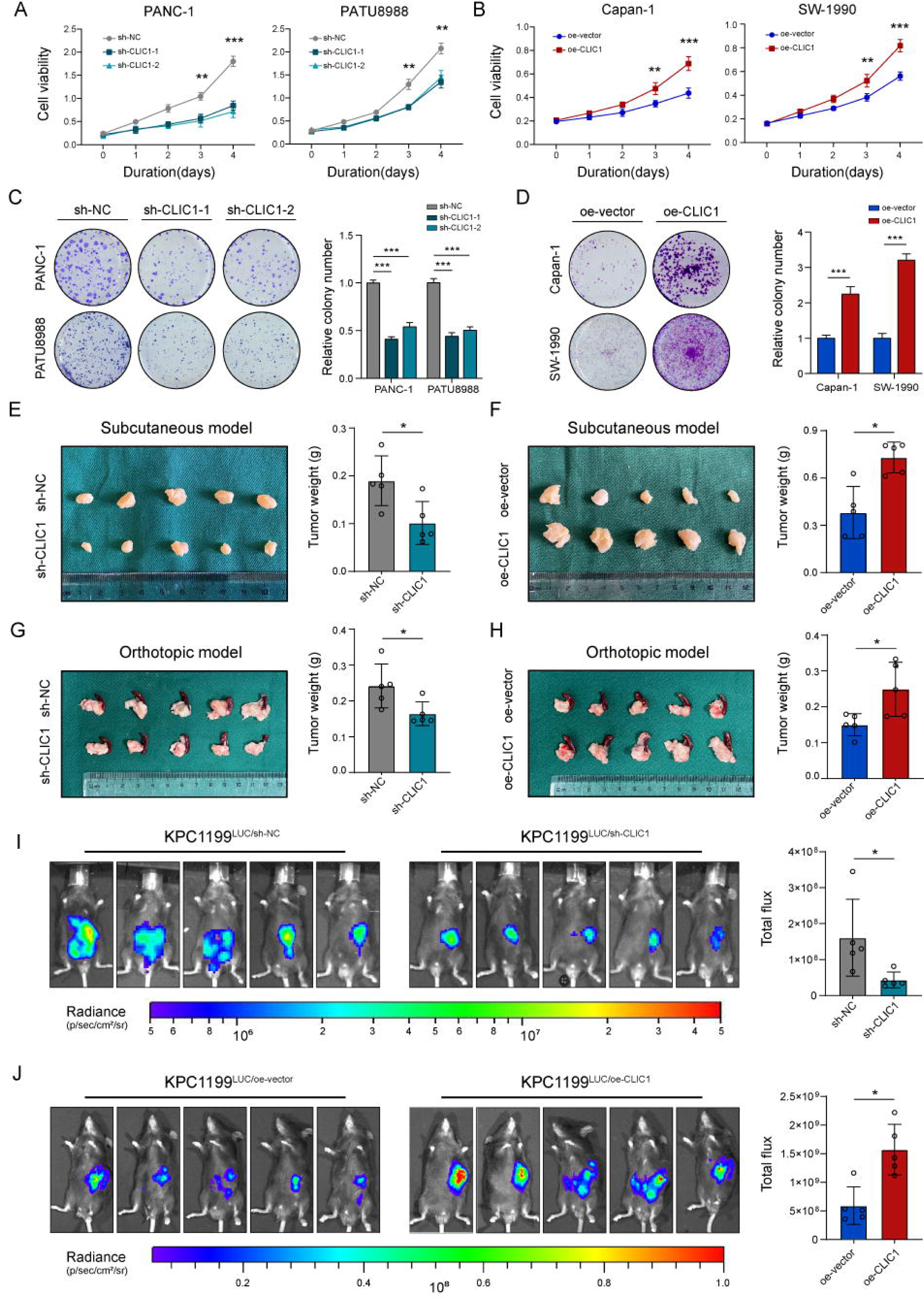
CLIC1 promotes the proliferative capacity of PDAC cells *in vivo* and *in vitro.* (A-B) The proliferation ability of PANC-1 and PATU8988 cells transfected with shRNA and Capan-1 and SW1990 cells overexpressing CLIC1 detected by CCK-8 assay (mean ± SEM., two-way ANOVA with Sidak’s multiple comparison test). (C-D) The colony formation ability of PANC-1 and PATU8988 cells with CLIC1 knockdown and Capan-1 and SW1990 cells with CLIC1 overexpression (mean ± SEM., two-tailed unpaired *t* test). (E-F) Tumor formation of CLIC1*^KD^* and CLIC1*^OE^* cells in nude mouse subcutaneous xenotransplantation models (n = 5 mice per group, mean ± SEM., Mann-Whitney U test). (G-H) Growth of orthotopic PDAC xenografts from CLIC1*^KD^* and CLIC1*^OE^* cells in C57BL/6 mice (n = 5 mice per group, mean ± SEM., Mann-Whitney U test). (I-J) Representative bioluminescence images and quantification of orthotopic C57BL/6 mice of the CLIC1*^KD^* and CLIC1*^OE^* cells and their control groups on day 15 (n = 5 mice per group, mean ± SEM., Mann-Whitney U test). \**P* < 0.05, \*\**P* < 0.01, \*\*\**P* < 0.001.

To explore whether the results of *in-vitro* experiments can be recapitulated *in vivo*, we established subcutaneous and orthotopic xenograft models. Compared to xenografts from the control group, mice injected with CLIC1-knockdown (CLIC1*^KD^*) cells exhibited a lower tumor burden, and correspondingly, a higher tumor burden was observed in mice injected with CLIC1-overexpressing (CLIC1*^OE^*) cells (Figure 5E-F). Meanwhile, the same trend was observed in the orthotopic PDAC model, which was built in mice by injecting CLIC1*^KD^* and CLIC1*^OE^* KPC*^luc^* cells into the pancreas of C57BL/6 mice (Figure 5G-H). In addition to measuring tumor weight, we also monitored orthotopic tumor growth by bioluminescence imaging and luminescence intensity, and immunohistochemistry for PCNA and Ki67. Compared to the control group, the CLIC1*^OE^* mice had a higher fluorescence flux (Figure 5J), whereas the opposite was true for the CLIC1*^KD^* group (Figure 5I). Concurrently, the IHC results of the specimens also indicated a stronger proliferative capacity in the control group than in the CLIC1*^KD^* group (Supplementary Figure 3G-H), with the opposite trend observed in the CLIC1*^OE^* group (Supplementary Figure 3I-J). Overall, these results suggest that CLIC1 confers a growth advantage to PDAC cells *in vivo*.

### CLIC1 promotes the growth of PDAC cells by enhancing the Warburg effect

To determine the mechanism by which CLIC1 promotes PDAC progression, we conducted GSEA using the GEO and TCGA public databases. The HALLMARK_GLYCOLYSIS pathway in multiple databases was enriched in samples with high *CLIC1* expression (Supplementary Figure 4A). This result was also supported by the gene correlation analysis based on the TCGA cohort, which showed a significant positive correlation between the expression of *CLIC1* and two commonly used glycolytic signatures (Supplementary Figure 4B). These findings prompted us to speculate that CLIC1 may play a role in regulating aerobic glycolysis in PDAC. The idea that CLIC1 regulates the Warburg effect in PDAC cells is supported by the following results. (I) The qPCR results showed that the genetic manipulation of CLIC1 robustly affected the mRNA expression of glucose transporters and many glycolysis components (Supplementary Figure 4C-D). (II) CLIC1 knockdown impaired the ECAR (an indicator of glycolysis), whereas CLIC1 overexpression had the opposite effect (Supplementary Figure 4E). (III) Glucose uptake and lactate production were enhanced in CLIC1*^OE^* cells, and CLIC1*^KD^* cells showed the opposite result (Supplementary Figure 4F-G). (Ⅳ) In IHC staining of 14 PDAC samples that had undergone 18F-fluorodeoxyglucose (18F-FDG) PET-CT, high CLIC1 expression was accompanied by a higher SUV-max, reflecting metabolic activity, in contrast to low expression (Supplementary Figure 4H-I). Furthermore, the addition of two glycolytic inhibitors (2-DG and galactose) to the colony formation model significantly inhibited the ability of CLIC1 to promote colony formation, inferring that CLIC1 promotes cell proliferation via aerobic glycolysis (Supplementary Figure 4J-K).

### CLIC1 maintains the stabilization of HIF1_α_

To further investigate the potential mechanisms by which increased CLIC1 expression contributes to the Warburg effect, GSEA based on transcriptome data from CLIC1*^KD^* cells and their control counterparts was conducted. The HALLMARK_HYPOXIA gene set was significantly enriched in the CLIC1 high-expression group, which was supported by GEO data (Figure 6A). HIF1α is a critical regulator of the Warburg effect^27^, and given that multiple glycolytic enzymes regulated by CLIC1 are important transcriptional targets of HIF1α, we hypothesized that CLIC1 participates in aerobic glycolysis by modulating HIF1α. Furthermore, the mRNA levels of *HIF1*_α_ were not affected by *CLIC1*, indicating that CLIC1-mediated regulation of HIF1α is either translational or posttranslational (Figure 6B).

**Figure 6.**
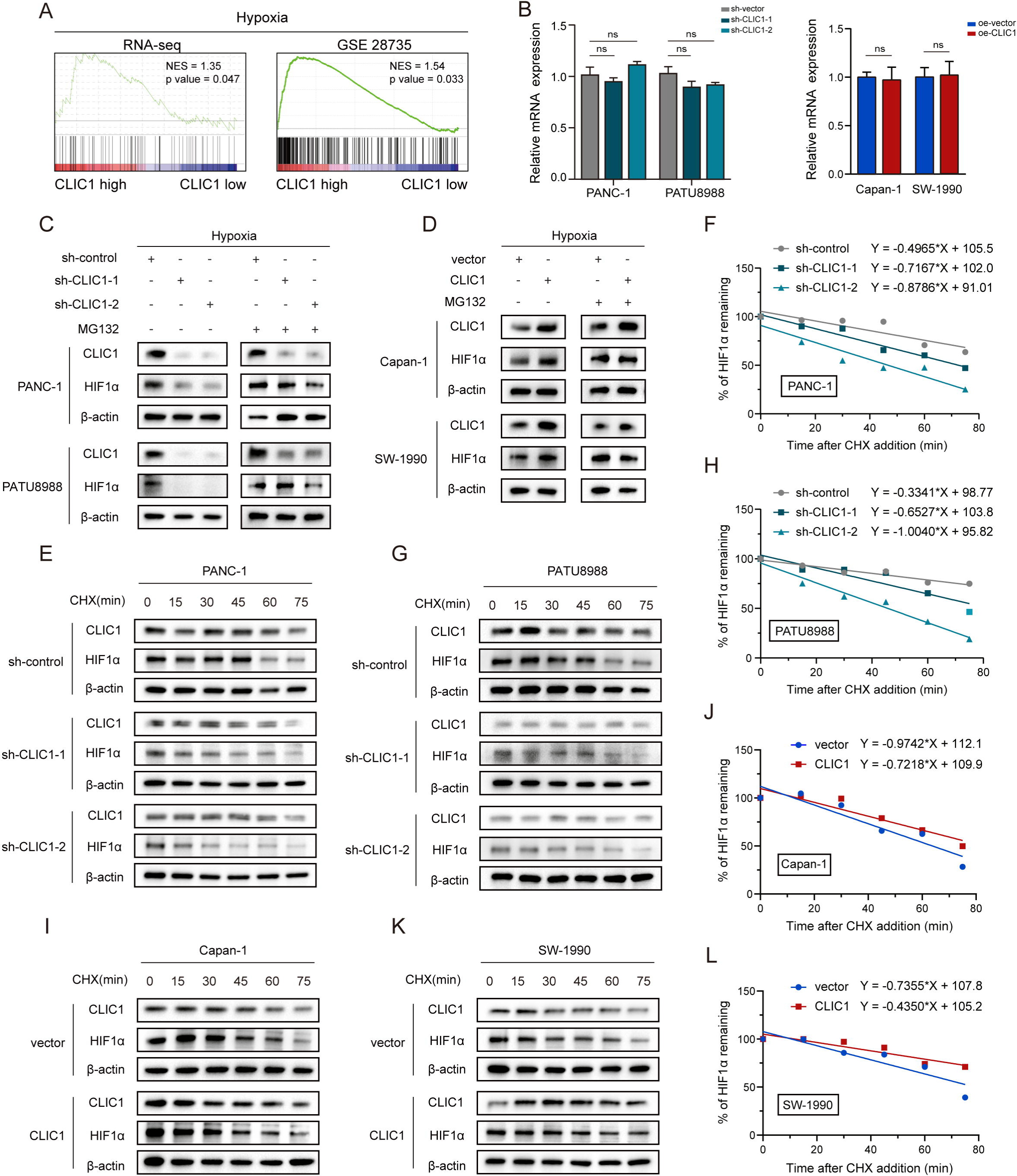
CLIC1 maintains the stabilization of HIF1α. (A) GSEA of *CLIC1* expression in the hypoxia gene set based on RNAseq and GEO data. (B) Expression of *HIF1*_α_ under hypoxic conditions in CLCI1*^KD^* and CLCI1*^OE^* and their control cells (mean ± SEM., two-tailed unpaired *t* test). (C-D) Altered expression of HIF1α protein levels in CLCI1 knockdown and overexpression and their control cells treated with or without 10 μM MG132 for 6 h under hypoxia. (E-H) Determination of HIF1α stability in CLCI1*^KD^* and control cells incubated with 20 μg/mL cycloheximide (CHX) for the indicated times after 6 h of hypoxia. (I-L) Stability assay of HIF1α in CLIC1*^OE^* and vector control cells exposed to hypoxia for 6 h followed by incubation with 20 μg/mL CHX until the specified time. ns, No significance.

As expected, under hypoxic conditions, HIF1α protein levels were significantly reduced after CLIC1 knockdown (Figure 6C) and increased after CLIC1 overexpression (Figure 6D). It is widely recognized that hydroxylation-modified HIF-1a protein readily binds to the specific E3 ubiquitin ligase Von HippelLLindau (VHL), which leads to degradation via the ubiquitinLproteasome pathway and is the main factor for the extreme instability of HIF-1a protein under normoxia^28^. Treatment with MG132, a proteasome inhibitor, eliminated the alterations in HIF1α expression caused by CLIC1 loss-of-function (Figure 6C) or gain-of-function (Figure 6D). These results suggest that CLIC1 regulates the posttranslational stability of the HIF1α protein and that this regulatory mechanism is mediated by the proteasomal pathway. Next, cells were exposed to hypoxia to stabilize HIF1α protein and then treated with cycloheximide (CHX) to block protein synthesis. Notably, CLIC1 knockdown shortened the half-life of HIF1α under hypoxic conditions (Figure 6E-H), while CLIC1 overexpression resulted in slower degradation of the HIF1α protein (Figure 6I-L), indicating that CLIC1 stabilizes HIF1α.

### CLIC1 suppresses ROS-dependent hydroxylation of HIF1_α_

We then explored how CLIC1 regulates the stability of the HIF1α protein. Given that many oncogenes have been shown to stabilize the HIF1α protein by inhibiting hydroxylation^29–31^, we tested whether CLIC1 stabilizes HIF1α by affecting its proline hydroxylation. To measure the hydroxylated HIF1α levels, cells were pretreated with MG132 to prevent hydroxylated HIF1α from degradation. In this process, less HIF1α accumulated in CLIC1*^KD^* cells, but significantly more hydroxylated HIF1α accumulated (Figure 7A), whereas the opposite was observed in CLIC1*^OE^* cells (Figure 7B), indicating low HIF1α hydroxylation in the presence of CLIC1.

**Figure 7.**
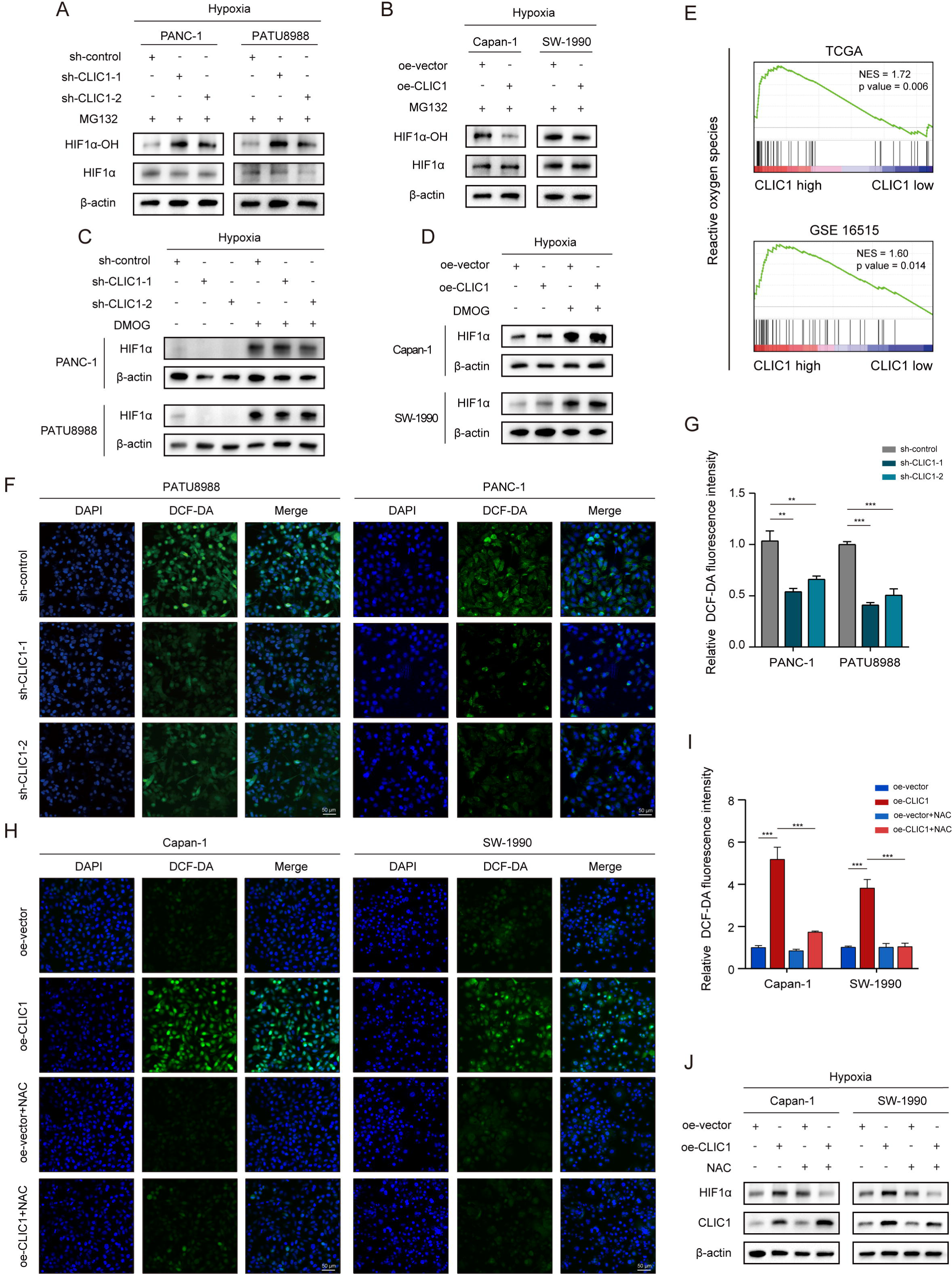
CLIC1 suppresses ROS-dependent hydroxylation of HIF1α. (A-B) Expression levels of HIF1α-OH and HIF1α in CLIC1 knockdown and overexpression and control cells treated with 10 μM MG132 for 6 h under hypoxia. (C-D) HIF1α expression in CLCI1*^KD^* and CLCI1*^OE^*and their control cells treated with or without 1 mM dimethyloxalylglycine (DMOG) for 24 h under hypoxic conditions. (E) Based on TCGA and GEO data, GSEA of CLIC1 expression in reactive oxygen species (ROS) gene set enrichment. (F-G) Fluorescence imaging and quantitative analysis of the ROS levels in CLIC1 knockdown and control cells under hypoxic conditions (mean ± SEM., two-tailed unpaired *t* test, scale bar: 50 μm). (H-I) Fluorescence imaging and quantitative analysis of the ROS levels in CLIC1*^OE^* and control cells treated with or without 10 mM N-acetylcysteine (NAC) for 24 h under hypoxia (mean ± SEM., one-way ANOVA with Tukey’s multiple comparison test, scale bar: 50 μm). (J) Expression of HIF1α in CLIC1-overexpressing and vector control cells treated with 10 mM NAC for 24 h under hypoxic conditions. \*\**P* < 0.01, \*\*\**P* < 0.001.

Prolyl hydroxylase domain-containing proteins (PHDs) can hydroxylate HIF1α in an oxygen-dependent manner at two proline sites (Pro-402 and Pro-564)^32^. Therefore, we further investigated whether CLIC1 regulates HIF1α hydroxylation through a PHD-dependent mechanism. This was verified by measuring the effect of dimethyloxalylglycine (DMOG), a potent PHD inhibitor, on HIF1α protein stability. If CLIC1 affects HIF1α stability by regulating PHD activity, DMOG treatment would abolish the effect of CLIC1 expression regulation. Equal levels of HIF1α were indeed observed in CLIC1*^KD^*cells (Figure 7C) and CLIC1*^OE^* cells as well as in their control cells (Figure 7D) in response to DMOG treatment. Overall, these results suggest that CLIC1 reduces PHD activity to hinder hydroxylation and enhance HIF1α protein stability.

In addition to intracellular oxygen concentration, the activity of PHDs can be regulated by numerous intracellular signals including reactive oxygen species (ROS), which have been confirmed to inhibit PHDs to stabilize HIF1α^33^. It has been reported that CLIC1 can act as a sensor and effector in the oxidative stress state of cells. In other words, CLIC1 is anchored to the plasma membrane after altering its structure by releasing GSH, which allows sustained ROS production by promoting an increase in chloride permeability^34^. Therefore, we infer that CLIC1-mediated HIF1α stabilization is contingent on ROS production. GSEA was performed using the TCGA and GEO databases and showed positive correlations between CLIC1 expression and activation of oxidative stress-related signaling pathways (Figure 7E). To test whether CLIC1 affected the expression of HIF1α via ROS, we first examined the effect of CLIC1 expression on intracellular ROS levels with an ROS assay kit. ROS levels were estimated with 2,7-dichlorofluorescein diacetate (DCF-DA). The fluorescence intensity of DCF-DA was weaker in CLIC1*^KD^* cells than in control cells (Figure 7F-G) but was inversely stronger in CLIC1*^OE^* cells and significantly reduced after the application of the antioxidant N-acetylcysteine (NAC) (Figure 7H-I), suggesting that CLIC1 upregulated intracellular ROS production. Interestingly, HIF1α levels were no longer increased in CLIC1*^OE^* PDAC cells after NAC was applied, indicating that NAC inhibits the oxidative stress state, thereby hijacking the downstream effects mediated by the CLIC1-ROS axis (Figure 7J).

## DISCUSSION

The insight that mechanical forces regulate cellular function permeates several areas of research, including metabolism. In turn, cellular metabolism has broad potential to regulate a large variety of cellular processes^12^. Understanding how cellular metabolism is regulated by the physical microenvironment and how alterations in tissue mechanics contribute to tumor development is a promising topic^35^. Based on the typical pathological features of PDAC, the mesenchymal-rich tumor microenvironment, this study focuses on the two major issues of the tumor matrix and metabolic reprogramming in PDAC and innovatively proposes that CLIC1 associates matrix stiffness with the Warburg effect to promote tumor growth.

Cells can sense mechanical signals from the ECM via integrins^36^, which further form integrin-adhesion complexes containing proteins involved in structural support and signal transduction, resulting in coupling of the mechanical environment to intracellular signaling pathways^37^. The transcription factors YAP and TAZ have been identified as the primary mechanotransduction factors apart from integrins and their downstream effectors^25^. Despite increasing evidence linking matrix stiffness, integrins, and YAP/TAZ, the connection between Wnt/β-catenin signaling and ECM stiffness remains poorly understood. However, our data suggest that PDAC cells can modulate Wnt/β-catenin/TCF4 signaling in response to ECM stiffness. This conclusion is also supported by Tao B et al., who found that matrix stiffness promotes glioma stemness through activation of the BCL9L/Wnt/β-catenin signaling pathway^38^.

Based on previous research, CLIC1 is upregulated in various malignant tumors, and its expression is associated with poor outcomes^19–21^. Aberrant CLIC1 expression endows tumor cells with high motility and invasiveness, making it a significant contributor to tumor invasion, metastasis, and angiogenesis^39–41^. In our study, we found that CLIC1 plays a critical role in cell-extracellular matrix interactions but differs in that it is a key molecule linking matrix stiffness and glycolysis. We discovered that PDAC cells can sense extracellular matrix stiffness and upregulate CLIC1 expression through the Wnt/β-Catenin/TCF4 signaling pathway, which promotes tumor growth by enhancing glycolytic capacity through ROS/HIF1α signaling (Figure 8). This will provide new insights into the biological characteristics of PDAC and offer a new perspective for its treatment.

**Figure 8.**
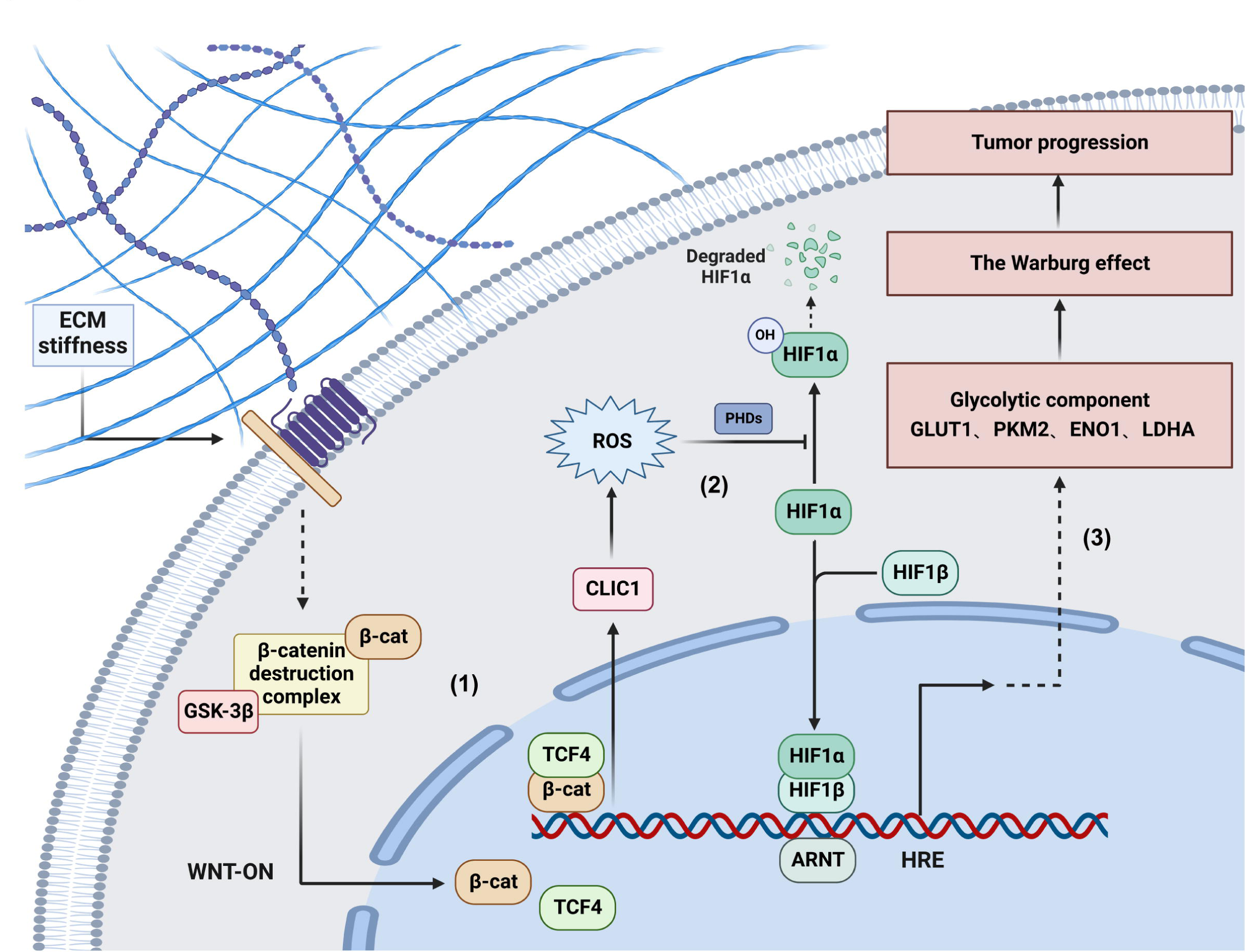
Model diagram of the regulatory mechanism of CLIC1 in PDAC progression. (1) PDAC cells sense ECM stiffness and upregulate CLIC1 expression through the Wnt/β-Catenin/TCF4 signaling pathway. (2) CLIC1 inhibits the hydroxylation of HIF1α to stabilize it by promoting the production of ROS. (3) CLIC1 drives PDAC progression through the Warburg effect.

In recent years, the concept of “new uses for old drugs” has received considerable attention in anticancer therapy. Metformin, a potent inhibitor of CLIC1 activity, suppresses the growth of glioblastoma stem cells by blocking their ion channel function^42^. The functional expression of CLIC1 determines the sensitivity of cancer cells to biguanides, which is necessary for the antitumor effect of biguanides^43, 44^. Moreover, metformin has been reported to exert its anticancer effect by directly or indirectly inhibiting the Warburg effect. For example, metformin promotes BG45-induced apoptosis through the anti-Warburg effect in cholangiocarcinoma cells^45^. The combination of 2-DG and metformin synergistically inhibits the energy metabolism of cancer cells and reduces ATP levels^46^. This combination therapy is more effective in drug-resistant pancreatic cancer cells with defective glycolysis, indicating that metformin sensitizes cancer cells to 2-DG^47^. Therefore, from the perspective of precision medicine, the identification of CLIC1-positive tumors, combined with PET-CT for the assessment of glycolytic metabolism, may allow for the selection of patients with specific types of tumors who could benefit from metformin treatment, thereby enhancing drug activity and reducing systemic toxicity.

In conclusion, our research revealed a previously unprecedented role of CLIC1, wherein increased CLIC1 expression induced by matrix stiffness promotes the Warburg effect in PDAC. Moreover, our findings provide new mechanistic insights into the key effects of CLIC1 in HIF1α stabilization by impairing its hydroxylation in a ROS level-dependent manner. The above findings suggest that targeting the CLIC1/ROS/HIF1α axis to inhibit aerobic glycolysis and reverse the malignant progression of PDAC is highly feasible.

## Abbreviations

18F-FDG: 18F-fluorodeoxyglucose
AnimalTFDB: animal transcription factor database
C-ECM: cancer-associated ECM
CHX: cycloheximide
CLIC: chloride intracellular channel
CLIC1: chloride intracellular channel 1
DCF-DA: 2,7-dichlorofluorescein diacetate
DEG: differentially expressed genes
DMOG: dimethyloxalylglycine
ECM: extracellular matrix
GSVA: gene set variation analysis
IHC: immunohistochemistry
KPC: *Kras*^G12D^/*Trp53*^R172H^/*Pdx1*-Cre
NAC: N-acetylcysteine
PanINs: pancreatic intraepithelial neoplasms
PDAC: pancreatic ductal adenocarcinoma
PET-CT: positron emission tomography-computed tomography
PHDs: prolyl hydroxylase domain-containing proteins
PSR: picrosirius red
ROS: reactive oxygen species
SEM: standard error of the mean
SUV-max: standardized uptake value-max
TMA: tumor micro array
TME: tumor microenvironment
VHL: Von HippelLLindau

## Author contributions

D.-J.L., S.-H.J., P.-X.J. and J.-H.Z. were responsible for concept and experimental design; Y.-W.S., W.L. and Y.-M.H. were responsible for clinical samples collection; Q.L., L.-P.H., J.Y., H.-F.Y. and Y.-W.S. were responsible for animal experiments; Z.-H.D., Y.-H.Z., P.-X.J., J.-H.Z. and R.-K.Z. performed cell molecular biology experiments; P.-X.J., Y.-H.Z., R.-K.Z. and W.L. were responsible for data measurement and scientific writing; D.-J.L., Q.-Y.J. and Y.-F.Y. were responsible for paper revision.

## Acknowledgments

We thank Dr. Lin-Li Yao, Xiao Li, Shan Zhang for the technical supports.

