## Supplemental Material for "A CLIC1 network coordinates matrix stiffness and the Warburg effect to promote tumor growth in pancreatic cancer"

**Supplementary Experimental Procedures**

**Cell culture and reagents**

The human PDAC cell lines AsPC-1, BxPC-3, Capan-1, CFPAC-1, MIA PaCa-2, PANC-1, PATU8988, and SW1990 and the murine PDAC cell line KPC1199 used in this study were all preserved in Shanghai Cancer Institute, Ren Ji Hospital, School of Medicine, Shanghai Jiao Tong University. All cells were cultured in the recommended medium according to American Type Culture Collection protocols supplemented with 10% fetal bovine serum (FBS) and 1% streptomycin/penicillin (P/S) in a humidified incubator under 5% $CO_2$ at 37 °C. For studies on hypoxia, PDAC cells were grown in an atmosphere replacing oxygen with $N_2$ in a HERAcell 150i $CO_2$ incubator (Thermo Scientific, USA). The reagents used in the study were as follows: D-galactose (Sigma–Aldrich, #G5388), 2-DG (Sigma–Aldrich, #D8375), CHX (Sigma–Aldrich, #239763-M), DMOG (Selleck, #S7483), MG132 (Selleck, #S2619), NAC (Sigma–Aldrich, #A7250), PF-573228 (AbMole, #M2931), RGDS (AbMole, #M13751) and ICG-001(AbMole, #M2008). As previously described[1], to perform *in vitro* cell experiments simulating different matrix stiffnesses, prefabricated 6-well and 12-well cell culture plates with 0.5 kPa and 12 kPa hydrogels bound with type I collagen from bovine skin or rat tail (Matrigen, SW6-COL-0.5, SW6-COL-12, SW12-COL-0.5, SW12-COL-12) were used for cell seeding and culture.

**Immunohistochemistry (IHC)**

IHC was performed as previously described[2]. The primary antibody used for IHC staining were: collagen type I (14695-1-AP, 1:200, Proteintech), HK2 (66974-1-Ig, 1:200, Proteintech), GLUT1 (21829-1-AP, 1:200, Proteintech), LDHA (19987-1-AP, 1:200, Proteintech), and CLIC1 (sc-81873, 1:200, Santa Cruz). Positive scores were based on the percentage of positively stained cells (0 = 0-5%, 1 = 6-35%, 2 = 36-75% and 3 = 76-100%) and staining intensity (0 = no staining, 1 = weak staining, 2 = moderate staining, and 3 = strong staining). Final scores were obtained by multiplying the two values described above ("-" for a score of 0-1, "+" for a score of 2-3, "++" for a score of 4-6, and "+++" for a score of 7-9). Low expression was defined as a total score < 4, whereas high expression was correspondingly described as a total score ≥ 4.

### Picrosirius red (PSR) staining

Sirius Red (Servicebio, G1018) was used to prepare the PSR solution. Paraffin sections were dewaxed and incubated with xylene for 5 min twice, followed by 100%, 100%, and 75% ethanol for 5 min each. Then, the sections were rinsed under running tap water for 1 min. Subsequently, PSR staining was performed on the sections for 8 min, and then the sections were dehydrated quickly with anhydrous ethanol three times. Finally, the sections were incubated in 100% ethanol and xylene for 5 min 3 times each before sealing with neutral gum. Microscope inspection (NIKON ECLIPSEE 100), image acquisition (NIKON DS-U3) and analysis were then conducted.

### Quantitative real-time PCR (qRT–PCR)

Total RNA of PDAC cells was extracted using RNAiso Plus (Takara Bio, #9109) and reverse transcribed into cDNA by the PrimeScript$^{TM}$ RT Reagent Kit (Takara Bio, #RR037A). Afterwards, the real-time PCR assay was performed with TB Green Premix Ex Taq$^{TM}$ II (Takara Bio, #RR820A) on an Applied Biosystems$^{TM}$ 7500 Real-Time PCR System (Thermo Scientific, USA) according to the recommended thermal cycle settings (a 10 min initial cycle at 95.0 °C, then 40 cycles of 10 s at 95.0 °C to 30 s at 60.0 °C). Relative mRNA expression was normalized to the *18S* RNA level. The primer sequences are listed in Supplementary Table 1.

### Western blotting analysis

In this report, a modified protocol based on the method reported by R. Li et al. was used[3]. The antibodies used in this study were anti-CLIC1 (sc-81873, 1:1000, Santa Cruz), anti-HIF1α (ab2185, 1:1000, Abcam), anti-HIF1α-OH (#3434, 1:1000, Cell Signaling Technology), anti-TCF4 (22337-1-AP, 1:1000, Proteintech), anti-SPI1 (55100-1-AP, 1:1000, Proteintech), anti-FOXA2 (22474-1-AP, 1:1000, Proteintech), anti-EGR2 (13491-1-AP, 1:1000, Proteintech), anti-β-actin (ab8226,1:5000, Abcam) and anti-Histone-H3 (17168-1-AP, 1:5000, Proteintech). The species-specific secondary antibodies included in this report are as follows: HRP-conjugated goat anti-rabbit IgG (GB23303, 1:8000, Servicebio) and HRP-conjugated goat anti-mouse IgG (GB23301, 1:8000, Servicebio). The bound secondary antibodies were detected by a chemiluminescence fluorescence imaging system (BIO-RAD, ChemiDoc XRS+, 1708265).

**Knockdown and overexpression assay**

The plasmid overexpressing CLIC1 (NM_001288) was constructed by GeneChem (Shanghai, China). The short hairpin RNA (shRNA) sequences targeting the CLIC1 gene were as follows: sh-1, GAGCUUGUGUUGCUGAATT and sh-2, GCCAAAGUUACACAUAGUATT. The lentiviral vector hU6-MCS-Ubiquitin-EGFP-IRES-puromycin was used for CLIC1 overexpression and knockdown assays. For lentiviral transfection, cells were plated 24 h in advance and a lentiviral suspension was added to the medium in the presence of 25× HitransG P (GeneChem, Shanghai, China). After 48 h of transfection, 5 μg/mL puromycin was added to the medium to screen for stable cell lines for 2 weeks. For small interfering RNA (siRNA) transfection, Lipofectamine RNAiMAX (Thermo Scientific, #13778150) was used to transfect specific siRNA into PDAC cells to knock down gene expression. Following the manufacturer 's instructions, cells transfected with siRNA were collected after 48 h for qRT–PCR or western blotting to detect knockdown efficiency or further assays. The siRNA oligonucleotides were synthesized by Obio (Shanghai, China). The detailed sequences of the siRNAs used in this study are as follows: si-TCF4-1, 5′-GGCCUCAUCGUCUCCUAAUUATT-3′; si-TCF4-2, 5′-UCCGAGAAAGGAAUCUGAAUCTT-3′; si-FOXA2-1, 5′-GGGAUGAACGGCAUGAACACGUACA-3′ and si-FOXA2-2, 5′-GGAUGAACGGCAUGAACACGUACAU-3′;

**Cell proliferation assays**

Cell proliferation was detected by the Cell Counting Kit-8 (CCK-8, MCE, #HY-K0301) assay. A total of 2000 cells were seeded into 96-well plates with quintuple samples each. After cell adherence, the culture medium was replaced with 10 % CCK-8 medium. The 96-well plates were incubated at 37 °C for 2 h, and then the absorbance values at 450 nm were measured using an automatic enzyme-linked immune detector (Bio-Rad Laboratories, Hercules, CA). Measurements were recorded every 24 h for 4-5 days, and data were collected and plotted as growth curves for comparative analysis. The experiments were repeated three times independently. For the colony formation assessment, cells were inoculated in 6-well plates at a ratio of 500-1000/well and cultured for 12-14 days. Then, the colonies were fixed using 4% paraformaldehyde (Servicebio, #G1101) for 30 minutes and stained with 0.1% crystal violet (Servicebio, #G1014) for another 30 minutes. Then, photographs were taken, and colonies were counted. Colonies of more than 30 cells were counted under a microscope. Three independent experiments were performed.

**Measurement of ROS levels**

The ROS fluorescent probe dihydroethidium (ShareBio, 38483-26-0) was used to measure the ROS levels of the cells according to the manufacturer's protocol. The indicated PATU8988 cells to be measured were seeded into chambered coverslips (Ibidi, 80826) in DMEM containing 10% FBS and incubated at 37 °C with 5% $CO_2$ for 48 h. Then, we removed the medium and rinsed the chambers with PBS. Dihydroethidium (DHE) was diluted in serum-free medium at 1:1000 to a final concentration of 10 μmol/L, and then diluted DHE was added to each chamber at a volume of 170 μL. After a 20-min incubation at 37 °C in the cell incubator, the cells were washed three times with PBS to fully remove the DHE that did not enter the cells. The cell samples loaded with probes in situ were observed directly by laser confocal microscopy (Leica, Germany) with an excitation wavelength of 488 nm and emission wavelength of 525 nm. The relative 2,7-dichlorofluorescein diacetate (DCF-DA) was calculated in 3 randomly selected regions under 400× magnification in triplicate per sample. The experiments were repeated three times.

**Seahorse analyses**

A Seahorse XF96 Flux Analyzer (Seahorse Bioscience, Agilent) was used to determine the extracellular acidification rate (ECAR) of the cells. Briefly, the cells were seeded at a density of $1 \times 10^4$ per well in XF96-well plates and processed as specified according to the manufacturer's instructions. Prior to the assay, the culture medium was replaced with the assay medium. ECAR were measured using the Glycolysis Stress Test Kit (Seahorse, #103020-100) and the Mitochondrial Stress Test Kit (Seahorse, #103015-100), respectively. Both measurements were normalized by total protein content. This experiment was performed in quintuplicate per sample three times independently.

**Glucose and lactate measurements**

According to the manufacturer’s instructions, cells were cultured in six-well plates for 24 h and the supernatant was collected for further detection. To investigate the changes in glucose consumption and lactate production in cells, the Glucose Uptake Assay Kit (Abcam, #ab136955) and L-Lactate Assay Kit (Abcam, #ab65330) were used to determine the glucose and lactate contents of cells, respectively. The results were normalized to the total protein amount. This experiment was performed three times independently.

**Animal model studies**

The nu/nu and C57BL/6 mice were purchased from East China Normal University, whereas the Pdx1-Cre, LSL-Kras$^{G12D/+}$, and LSL-Trp53$^{R172H/+}$ mice were purchased from Jackson Laboratory. The mice were manipulated and housed according to the criteria in the Guide for the Care and Use of Laboratory Animals of the National Institutes of Health (Bethesda, MD). All manipulations were performed following the approved protocol number 20141204, assigned by the Research Ethics Committee of East China Normal University. We ensured that the tumor size/burden did not exceed 200 mm$^3$ throughout our study, as per the guidelines of the institutional review board. The investigators who conducted the animal experiments were blinded to allocation during the experiments and outcome assessments.

**Subcutaneous and orthotopic xenograft model**

For the subcutaneous xenograft model, athymic male nu/nu mice aged 6 to 8 weeks were used in this study. In this model, 1 × 10$^6$ cells (Capan-1 cells infected with CLIC1/Vector, 1 × 10$^6$ PATU8988 cells transfected with shcontrol/shCLIC1-1) resuspended in 200 μL of PBS were injected subcutaneously into one side of the back of each nude mouse. After 4 weeks, the tumors were resected for imaging and weighing. In the orthotopic xenograft model, 1 × $10^5$ $CLCI1^{KD}$ or $CLIC1^{OE}$ $KPC1199^{luc}$ cells were resuspended in 20 μL of PBS and transplanted directly into the body of the pancreas of the mice (C57BL/6, male, 6-8 weeks). Bioluminescence imaging was performed after intraperitoneal injection of D-fluorescein in 5 randomly selected mice per group at weeks 2 and 4 after inoculation. Four weeks after implantation, the mice were sacrificed, and the tumors were removed, fixed in 4% paraformaldehyde, weighed, and photographed.

**Bioinformatics analysis**

Using the "correlation analysis" module of Gene Expression Profiling Interactive Analysis 2.0 (GEPIA2) (http://gepia2.cancer-pku.cn/#index), the correlation coefficients (R) and p-values between gene sets or genes were obtained, and a scatter plot with $\log_2$TPM values was generated. To annotating the biological roles of these tumor matrix- and glucose metabolism-correlated genes, KOBAS-i database (http://bioinfo.org/kobas) was used to test the statistical enrichment of DEGs in Kyoto Encyclopedia of Genes and Genomes (KEGG) pathways. For gene set enrichment analysis (GSEA), we used the median expression value of CLIC1 as the cutoff. Gene expression above the median was designated as "high expression", and below was defined as "low expression". GSEA was performed on the GSEA software (version 4.2.3), with statistical significance (false discovery rate, FDR) set at 0.25.

**RNA sequencing**

Total RNA was extracted using RNAiso Plus reagent (Takara Bio, #9109) following the manufacturer’s protocol. Then, RNA quality was determined by a 5300 Bioanalyzer (Agilent) and quantified using an ND-2000 (NanoDrop Technologies). Only high-quality RNA samples (OD260/280=1.8~2.2, OD260/230≥2.0, RIN≥6.5, 28S:18S≥1.0, >1 μg) were used to construct the sequencing library. The samples underwent a process of RNA purification, reverse transcription, library construction and sequencing, which was performed at Shanghai Majorbio Bio-pharm Biotechnology Co., Ltd. (Shanghai, China) according to the manufacturer’s protocols (Illumina, San Diego, CA). The CLIC1 RNA-seq transcriptome library was prepared following Illumina® Stranded mRNA Prep, Ligation from Illumina (San Diego, CA) by using 1 μg of the total RNA. First, messenger RNA was isolated according to the polyA selection method by oligo(dT) beads and then fragmented by fragmentation buffer. Second, double-stranded cDNA was synthesized with a SuperScript double-stranded cDNA synthesis kit (Invitrogen, CA) along with random hexamer primers (Illumina). Then the synthesized cDNA was subjected to end-repair, phosphorylation, and 'A' base addition according to Illumina's library construction protocol. Libraries were size-selected for cDNA target fragments of 300 bp on 2% Low Range Ultra Agarose followed by PCR amplification using Phusion DNA polymerase (NEB) for 15 PCR cycles. After quantification by Qubit 4.0, a paired-end RNA-seq library was sequenced with a NovaSeq 6000 sequencer (2×150 bp read length).

**Proteomic analysis**

We provided 8 liquid-nitrogen-frozen PDAC tissues resected from patients in Ren Ji Hospital and entrusted the Shanghai Bioprofile for the isobaric tags for relative and absolute quantitation (iTRAQ) proteomic experiment. The samples underwent tissue lysis, protein extraction, BCA quantification, SDS–PAGE electrophoresis, Coomassie Brilliant Blue staining, and enzymatic hydrolysis (300 μg per sample) for preparation. Peptides were labeled with TMT reagents according to the manufacturer's protocol (Thermo Scientific). The fractions were dried for nano-LC–MS/MS analysis. The labeled peptides of each group were mixed in equal amounts. The dried peptides were separated using the Pierce™ High-pH Reversed-Phase Peptide Fractionation Kit (Thermo Scientific). Finally, the samples were combined into 10 components. After drying, the peptides of each component were redissolved in 0.1% FA for LC–MS analysis. An appropriate amount of peptide was taken from each sample for chromatographic separation using the Easy nLC 1200 chromatographic system (Thermo Scientific). After separation, the peptides were analyzed by data-dependent acquisition (DDA) mass spectrometry using Q-Exactive HF-X mass spectrometry (Thermo Scientific). The resulting LC–MS/MS raw files were imported into the search engine Sequest HT of Proteome Discoverer software (version 2.4, Thermo Scientific) for the database search. The database used for searching was UniProt-Homo sapiens (Human) [9606]-202249-210712.fasta (downloaded at 2021.07.12, including 2022249 protein sequences) from https://www.uniprot.org/taxonomy/9606. The false discovery rate (FDR) taken for filtering and exclusion was ≤ 0.01.

**Chromatin immunoprecipitation (ChIP) assay**

ChIP assays were performed with a ChIP assay kit (Pierce Agarose ChIP Kit, Thermo Scientific) according to the manufacturer’s protocol. The prepared DNA–protein complexes immunoprecipitated with anti-TCF4 (22337-1-AP, Proteintech) or control IgG (Cell Signaling) from the sonicated cell lysates with Premix Taq (Cell Signaling) were quantified by qPCR (SYBR Green system). The sequences of the primers used in the ChIP process are listed in Supplementary Table 2.

**Luciferase reporter assay**

Briefly, a luciferase reporter assay was performed as follows: PATU8988 cells were cotransfected with CLIC1 promoter WT/MT plasmids, TCF4 vector/overexpressing plasmids and pRL-TK Renilla plasmid using Lipofectamine 2000 (Invitrogen, Thermo Scientific, Inc.) for further analysis. The Dual-Luciferase® Reporter Assay System (E1910, Promega, Madison, WI, USA) was used to analyze the luciferase activity according to the manufacturer's protocol. Luciferase activity was normalized for transfection efficiency to Renilla activity, and the results represent the mean ± standard deviation of triplicate samples. The wild-type and mutant CLIC1 promoter regions and primers, as well as those for TCF4, are listed in Supplementary Table 3. For the TOP/FOP flash assay, PATU8988 cells were transfected with TOP/FOP and Renilla plasmids in the 0.5 kPa and 12 kPa Matrigen Softwell 12-well cell culture plates described above. After 36 h of transfection, the cells were lysed with passive lysis buffer (E1910, Promega, Madison, WI, USA), and reporter gene expression was assessed using the Dual-Luciferase® Reporter Assay System (E1910, Promega, Madison, WI, USA). Luciferase activity was normalized for transfection efficiency to Renilla activity, and the results represent the mean ± standard deviation of triplicate samples.

### Supplementary Tables

**Supplementary Table 1.** The primer sequences for qRT-PCR assays

| Gene symbol | Forward sequence | Reverse sequence |
|---|---|---|
| CLIC1 | GAAATCAGGTGTCCCATTCCAG | TGGGGAGGTCGCTTCTCAA |
| HIF1α | GAACGTCGAAAAGAAAAGTCTCG | CCTTATCAAGATGCGAACTCACA |
| GLUT1 | ATTGGCTCCGGTATCGTCAAC | GCTCAGATAGGACATCCAGGGTA |
| HK2 | AGCCCTTTCTCCATCTCCTT | GCTTGCCTACTTCTTCACGG |
| LDHA | ATGGCAACTCTAAAGGATCAGC | CCAACCCCAACAACTGTAATCT |
| GPI1 | CAAGGACCGCTTCAACCACTT | CCAGGATGGGTGTGTTTGACC |
| PFKL | GGTGCCAAAGTCTTCCTCAT | GATGATGTTGGAGACGCTCA |
| ALDOA | AACTTTCCTCTGCCTAGCCC | GTACAGGCACAGTCGCAGAG |
| TPI1 | AGCTCATCGGCACTCTGAAC | CCACAGCAATCTTGGGATCT |
| PGK1 | TTGACGAGAACGCTCAGGTTG | ACGGCCCATTCCAAACAATTAG |
| PGAM2 | AGAAGCACCCCTACTACAACTC | TCTGGGGAACAATCTCCTCG |
| ENO1 | GCCGTGAACGAGAAGTCCTG | ACGCCTGAAGAGACTCGGT |
| PDK1 | CTGTGATACGGATCAGAAACCG | TCCACCAAACAATAAAGAGTGCT |
| PKM2 | ATGTCGAAGCCCCATAGTGAA | TGGGTGGTGAATCAATGTCCA |
| FOXA2 | GGAGCAGCTACTATGCAGAGC | CGTGTTCATGCCGTTCATCC |
| TCF4 | GAAAGCTGCGTGTCTGAAAA | CATCTGTCCCATGTGATTCG |
| 18S | ATCACCATTATGCAGAATCCACG | GACCTGGCTGTATTTTCCATCC |

**Supplementary Table 2.** The primers used in ChIP-PCR

| Gene symbol | Forward Primer | Reverse Primer |
|---|---|---|
| CLIC1 | AGGTTACTGAAAAGGCGGGA | GCACGTCTTATGTCTCTGCC |
| Negative control | TATCCCCTGTCCCCATTCCA | CCACTGGCCACTTTGTACAC |

**Supplementary Table 3.** Sequence of wild-type and mutated CLIC1 promotor regions

| Promoter type | Constructing sequences |
|---|---|
| Wild-type | AGAGGTATCTATCATTCATGCATTCGGCAACTATTTAGTGCCTATTTTGGGCCAGGGGTTG |
| | GACTCAAAGCGGTGAACATAATGAAGTCACTGCTCCATAAAGCTTATTTGTGCGTCTGTG |
| | TGTGTGTGTGTGTGTGTGTGTGTGTGTGTGTGTTGCGGGGGTGGGGTTATGAGGGA |
| | GAAAGAAAAAAAATATATATATATATATATATAATCCCTTTAAATGCACCATCCTCCCCAGCTT |
| | TGTTCCTAGTACCCTGAGATGGGGAGATTCCTCCCCAGCCCCCCAACCCAAGAAGTTAG |
| | GAAAGATGGTGGGGGTGAGATGACCTAGCTGTGCTAACCAGTAATTGGAAAATCCCCTC |
| | CAGCAAGAAAGGTGGGGGAACAGAGTTAAGGGCTGGGTTAATGGTTACCCCTGGCAAAT |
| | CTGTTGAGCAATGGAGCTATAGAAACTTGGAGGAGGTTGGTGACAGCTCTGGGGAGTGT |
| | CAGGGAGGGACCCACCTTCCAATCTGGGGTGTGAAGAGATTAGGGACTAGTTTACCAAG |
| | CCCAGGAAGGGGAGGGAGTGGAAAGAGAAGCCCAGAGAGGGAAAGAGGAGCTACTGA |
| | GATAGGAGAAATGCAGGGACAGGGAGGTGAGATGGAGGGAAACCTCTGTTGCAGGATT |
| | TGGGGGTACAGCCCTGGCTCTGCTGGATGGTTCCAGGAGAGGGCAGCGTTGCCTGTCA |
| | CCTGGTAAATTAAGGCACGATACCCTGTGGGTAGTCATGCCAGCCAGCCAAAGTCAATAT |
| | TGATATCAAGGCAGCTGTAGCTATAAAGCTGTAGGAGAGAAAAGAGAGGCCGGGAGAGG |
| | CTGCCAAACCTGTTTGATCTTCAAGCTGGCCCCATTTACCTAAGGCTTCTCTTGGACACA |
| | GACCAGTTGGAACAGGAGAGGACCCTGAGAAGTAGAGTTGTTTTGATCCCCTCCCCTCA |
| | ATGGGAAGGGGTCCTGTGTGCAGATTTTGAGGGTCACCATGAGGGAATTCACACCCACA |
| | CAGAGGCATGGAATCATACCCTTACAGGATGTTACAGTTCAAGGGAACGAACACCAAAGA |
| | TAGCCTTTCTGTCGGGAGGAGAGAAAAAGGCCCTCCCACAAGCAGGAGAAAACCCACA |
| | GAAGAGGAAGCACAAGAGGAAACCACCAACTGCTGGTTTCCAGCAGCACATCACGTCA |
| | CTCCACCCCACTCCTCCCCCAAGTCTCTCCCTTTCTTTAAGTTTCAAGCTCTGTTTTAGTT |
| | CTGTGTTCTTGCACTCCACCATGGTTTTCTGGAACTGTAGGTCTTTTGTTGGACAGAGGG |
| | TGATGAGGAGGATAGAGAAGGGATGGTTGGACAGGAGAGAAATTCAGGATATGGAGGCT |
| | GGAATTCTGGGTTTATTTTTTCAGCAGGTCATAAAGGTTTAATAGAAATCAAGGTTACTGG |
| | AGAGAGAGCTGCTCCTCCTATGTCCTCCCTGCTTATTTATTTAGGTGTCCCCAAGGACTC |
| | TACTCCCACTATTCTGTCTGATCTTTCTTTATCCCCATGACAGGACCAGCTAACAACACCC |
| | CTACCCCACCCCCTATACACATACTTCAGGATCGGGCCATGATAATCCCACCCCTCTGCC |

| | |
|---|---|
| | CCATCTCCAAGGCAACCTGTCAGTGAGACGAGGACAAAGGGCACAGGAAGGGGCCCCA<br>ATAGGAAACATAAGTGGAAGCACAAAGCTGACCAAGCTACAGGAACAGACCCCTCCCTG<br>CAACAAAGCCCCTTGCCTCGGCTTTATGTTTCCTTCGCAAAAGACTTCGTCATCTCCCTT<br>CCCATCCCTAACTCTACTTTCTTTTTCTTTTCTTCTTCTTCTTTTTTTTTTTTTTTTTTGAGA<br>TGGAGTTTCGCTCTTATTGCCCAGGCTGGAGTGCAATGGCACCATCTCAGCTCACTGCA<br>ACCTTCACCTCCCGGGTTCAATTGATTCTCCTGCCTCAGCCTCCCAAGTAGCTGGGATTA<br>CAGGTGCCCACCACCCCGCCTGGCAAATTTTTG |
| Mutation | AGAGGTATCTATCATTCATGCATTCGGCAACTATTTAGTGCCTATTTTGGGCCAGGGGTTG<br>GACTCAAAGCGGTGAACATAATGAAGTCACTGCTCCATAAAGCTTATTTGTGCGTCTGTG<br>TGTGTGTGTGTGTGTGTGTGTGTGTGTGTGTGTGTTGCGGGGGTGGGGTTATGAGGGA<br>GAAAGAAAAAAAATATATATATATATATATATAATCCCTTTAAATGCACCATCCTCCCCAGCTT<br>TGTTCCTAGTACCCTGAGATGGGGAGATTCCTCCCCAGCCCCCCAACCCAAGAAGTTAG<br>GAAAGATGGTGGGGGTGAGATGACCTAGCTGTGCTAACCAGTAATTGGAAAATCCCCTC<br>CAGCAAGAAAGGTGGGGGAACAGAGTTAAGGGCTGGGTTAATGGTTACCCCTGGCAAAT<br>CTGTTGAGCAATGGAGCTATAGAAACTTGGAGGAGGTTGGTGACAGCTCTGGGGAGTGT<br>CAGGGAGGGACCCACCTTCCAATCTGGGGTGTGAAGAGATTAGGGACTAGTTTACCAAG<br>CCCAGGAAGGGGAGGGAGTGGAAAGAGAAGCCCAGAGAGGGAAAGAGGAGCTACTGA<br>GATAGGAGAAATGCAGGGACAGGGAGGTGAGATGGAGGGAAACCTCTGTTGCAGGATT<br>TGGGGGTACAGCCCTGGCTCTGCTGGATGGTTCCAGGAGAGGGCAGCGTTGCCTGTCA<br>CCTGGTAAATTAAGGCACGATACCCTGTGGGTAGTCATGCCAGCCAGCCAAAGTCAATAT<br>TGATATCAAGGCAGCTGTAGCTATAAAGCTGTAGGAGAGAAAAGAGAGGCCGGGAGAGG<br>CTGCCAAACCTGTTTGATCTTCAAGCTGGCCCCATTTACCTAAGGCTTCTCTTGGACACA<br>GACCAGTTGGAACAGGAGAGGACCCTGAGAAGTAGAGTTGTTTTGATCCCCTCCCCTCA<br>ATGGGAAGGGGTCCTGTGTGCAGATTTTGAGGGTCACCATGAGGGAATTCACACCCACA<br>CAGAGGCATGGAATCATACCCTTACAGGATGTTACAGTTCAAGGGAACGAACACCAAAGA<br>TAGCCTTTCTGTCGGGAGGAGAGAAAAAGGCCCTCCCACAAGCAGGAGAAAACCCACA<br>GAAGAGGAAGCACAAGAGGAAACCACCAACTGCTGGTTTCCAGCAGCACATCACGTCA<br>CTCCACCCCACTCCTCCCCCAAGTCTCTCCCTTTCTTTAAGTTTCAAGCTCTGTTTTAGTT<br>CTGTGTTCTTGCACTCCACCATGGTTTTCTGGAACTGTAGGTCTTTTGTTGGACAGAGGG |

TGATGAGGAGGATAGAGAAGGGATGGTTGGACAGGAGAGAAATTCAGGATATGGAGGCT

GGAATTCTGGGTTTATTTTTTCAGCAGGTCATAAAGGTTTAATAGAAATCAAGGTTACTGG

AGAGAGAGCTGCTCCTCCTATGTCCTCCCTGCTTATTTATTTAGGTGTCCCCAAGGACTC

TACTCCCACTATTCTGTCTGATCTTTCTTTATCCCCATGACAGGACCAGCTAACAACACCC

CTACCCCACCCCCTATACACATACTTCAGGATCGGGCCATGATAATCCCACCCCTCTGCC

CCATCTCCAAGGCAACCTGTCAGTGAGACGAGGACAAAGGGCACAGGAAGGGGCCCCA

ATAGGAAACATAAGTGGAAGCACAAAGCTGACCAAGCTACAGGAACAGACCCCTCCCTG

CAACAAAGCCCCTTGCCTCGGCTTTATGTTTCCTTCGCAAAAGACTTCGTCATCTCCCTT

CCCATCCCTAACTCTACTTTCTTTTTCTTTTCTTCTTCTTTTTTTTTTTTTTTTTTTGAGA

TGGAGTTTCGCTCTTATTGCCCAGGCTGGAGTGCAATGGCACCATCTCAGCTCACTGCA

ACCTTCACCTCCCGGGTTCAATTGATTCTCCTGCCTCAGCCTCCCAAGTAGCTGGGAT**G**

**CACTTGTAA**CACCACCCCGCCTGGCAAATTTTTG

---

Note: Highlight area is the mutation sequence.

**Supplementary Table 4.** The top 200 tumor matrix-correlated genes in TCGA cohorts

| Gene symbol | logFC | adj.P-Val | Gene symbol | logFC | adj.P-Val |
|---|---|---|---|---|---|
| *GXYLT2* | 1.505361 | 5.42E-23 | *FBLN1* | 2.095937 | 1.83E-16 |
| *BNC2* | 1.000121 | 1.5E-22 | *TIMP3* | 1.81254 | 1.84E-16 |
| *COLEC12* | 1.874084 | 1.5E-22 | *SULF1* | 2.241726 | 2.52E-16 |
| *FAP* | 1.791975 | 3.1E-22 | *LHFPL6* | 1.316796 | 2.52E-16 |
| *DDR2* | 1.578528 | 3.1E-22 | *SPIRE2* | 1.378325 | 2.55E-16 |
| *ZNF521* | 1.257597 | 3.1E-22 | *VCAN* | 2.06411 | 3.31E-16 |
| *GLT8D2* | 1.526777 | 3.1E-22 | *CNN3* | 1.23564 | 5.24E-16 |
| *FBN1* | 2.142898 | 5.81E-22 | *TAGLN* | 1.761397 | 5.42E-16 |
| *PRRX1* | 1.878227 | 1.89E-21 | *ISM1* | 1.65047 | 5.42E-16 |
| *ADAM12* | 2.036922 | 2.4E-21 | *PCOLCE* | 1.492976 | 7.6E-16 |
| *MSRB3* | 1.541903 | 3.08E-21 | *TNFAIP6* | 1.620034 | 8.88E-16 |
| *CCDC80* | 2.349814 | 5.22E-21 | *EFEMP2* | 1.304839 | 1.03E-15 |
| *NID2* | 1.431358 | 6.23E-21 | *ADAMTS2* | 1.412123 | 1.12E-15 |
| *GAS1* | 2.213702 | 6.23E-21 | *HTRA1* | 1.50538 | 1.18E-15 |
| *POGLUT2* | 1.059667 | 7.89E-21 | *MN1* | 1.02071 | 1.27E-15 |
| *CLMP* | 1.749558 | 8.02E-21 | *GJA1* | 1.39477 | 1.38E-15 |
| *VSTM4* | 1.09675 | 1.36E-20 | *SH3PXD2B* | 1.271743 | 1.5E-15 |
| *RNF144A* | 1.285716 | 1.36E-20 | *SCUBE2* | 1.142474 | 1.69E-15 |
| *CTSK* | 1.960663 | 1.81E-20 | *RFLNB* | 1.178785 | 1.72E-15 |
| *PDGFRL* | 1.681833 | 1.84E-20 | *MOXD1* | 1.641426 | 1.74E-15 |
| *OLFML3* | 1.7803 | 2.34E-20 | *TIMP2* | 1.293149 | 1.97E-15 |
| *THY1* | 1.662941 | 3.27E-20 | *INHBA* | 1.879275 | 2.07E-15 |
| *P4HA3* | 1.356237 | 7.83E-20 | *PPP1R16A* | 1.157882 | 2.07E-15 |
| *PMP22* | 1.311379 | 1.05E-19 | *DAB2* | 1.124719 | 2.81E-15 |
| *SPARC* | 1.9792 | 1.1E-19 | *LRRC32* | 1.248204 | 2.81E-15 |
| *ECM2* | 1.302378 | 1.14E-19 | *IRAG1* | 1.198126 | 2.85E-15 |
| *PLXDC2* | 1.614841 | 1.35E-19 | *COL5A1* | 2.020512 | 2.91E-15 |
| *FKBP7* | 1.010309 | 1.41E-19 | *CXCL12* | 1.824771 | 2.97E-15 |
| *OLFML1* | 1.340504 | 1.45E-19 | *OMD* | 1.655703 | 2.97E-15 |
| *TCF4* | 1.043905 | 1.5E-19 | *ZEB1* | 1.096452 | 3E-15 |
| *CCN4* | 1.824897 | 1.97E-19 | *TENM3* | 1.025869 | 3.25E-15 |
| *FERMT2* | 1.273991 | 2.11E-19 | *VGLL3* | 1.075897 | 3.25E-15 |
| *CCDC8* | 1.298375 | 2.78E-19 | *LAMA2* | 1.460412 | 3.33E-15 |
| *FAM180A* | 1.095519 | 5.72E-19 | *GREM1* | 2.158123 | 3.33E-15 |
| *SPON1* | 2.061831 | 5.79E-19 | *MRC2* | 1.646219 | 3.43E-15 |
| *GASK1B* | 1.317509 | 8.17E-19 | *SMO* | 1.037115 | 3.61E-15 |
| *FBXL7* | 1.114812 | 1.01E-18 | *DIO2* | 1.268096 | 3.99E-15 |
| *CALD1* | 1.699527 | 1.09E-18 | *RHPN1* | 1.398322 | 3.99E-15 |
| *COL8A1* | 2.189392 | 1.35E-18 | *KIRREL1* | 1.258877 | 4.17E-15 |
| *LOX* | 1.811852 | 1.35E-18 | *PDGFRB* | 1.449656 | 4.23E-15 |
| *PODN* | 1.716881 | 1.35E-18 | *BOC* | 1.232489 | 4.93E-15 |

| | | | | | |
|---|---|---|---|---|---|
| *CPXM1* | 2.06895 | 1.54E-18 | *MSR1* | 1.422198 | 5E-15 |
| *AOC3* | 1.580161 | 1.55E-18 | *COL12A1* | 2.106063 | 5E-15 |
| *ALDH1L2* | 1.088302 | 2.11E-18 | *BASP1* | 1.527696 | 6.05E-15 |
| *NEXN* | 1.506788 | 2.11E-18 | *JCAD* | 1.001944 | 6.48E-15 |
| *COL6A3* | 2.349724 | 2.66E-18 | *FPR3* | 1.596091 | 6.48E-15 |
| *HMCN1* | 1.153806 | 2.71E-18 | *GPR68* | 1.199195 | 7.08E-15 |
| *CD248* | 1.726218 | 2.71E-18 | *PCDH18* | 1.019506 | 8.59E-15 |
| *COL5A2* | 2.223822 | 2.92E-18 | *LRP1* | 1.231186 | 8.59E-15 |
| *DCN* | 2.015353 | 3.55E-18 | *COL1A1* | 2.36889 | 8.85E-15 |
| *MFAP5* | 2.263522 | 3.55E-18 | *FOLR2* | 1.597008 | 9.17E-15 |
| *FBLN2* | 1.92295 | 4.58E-18 | *SFRP2* | 2.850036 | 1.05E-14 |
| *COPZ2* | 1.306439 | 4.58E-18 | *CTHRC1* | 2.064275 | 1.08E-14 |
| *FNDC1* | 2.207326 | 5.03E-18 | *LMCD1* | 1.086502 | 1.12E-14 |
| *PXDN* | 1.520765 | 5.14E-18 | *MFAP2* | 1.654976 | 1.36E-14 |
| *CDH11* | 1.7308 | 5.17E-18 | *MYL9* | 1.464821 | 1.38E-14 |
| *CLIC1* | 1.383195 | 5.58E-18 | *TNFSF4* | 1.034197 | 1.67E-14 |
| *GFPT2* | 1.725717 | 6.2E-18 | *MRGPRF* | 1.166532 | 1.72E-14 |
| *RECK* | 1.06304 | 7.24E-18 | *SRPX* | 1.576249 | 1.77E-14 |
| *ADAMTS12* | 1.648522 | 7.48E-18 | *SLIT2* | 1.194357 | 1.84E-14 |
| *CRISPLD2* | 1.732059 | 8.37E-18 | *ITGA11* | 1.676475 | 1.94E-14 |
| *DCHS1* | 1.090731 | 8.94E-18 | *TSTD1* | 1.024418 | 2.02E-14 |
| *OLFML2B* | 1.880961 | 1.08E-17 | *MRAS* | 1.012498 | 2.03E-14 |
| *FIBIN* | 1.800408 | 1.17E-17 | *C1QTNF3* | 1.473058 | 2.07E-14 |
| *SRPX2* | 1.395165 | 1.18E-17 | *LTBP1* | 1.425572 | 2.09E-14 |
| *PDGFRA* | 1.575154 | 1.55E-17 | *NID1* | 1.299184 | 2.15E-14 |
| *RAB31* | 1.66951 | 1.6E-17 | *NOX4* | 1.002161 | 2.27E-14 |
| *FSTL1* | 1.679163 | 1.78E-17 | *EMILIN1* | 1.62575 | 2.55E-14 |
| *LUM* | 2.25008 | 1.78E-17 | *MMP19* | 1.381189 | 2.55E-14 |
| *COL1A2* | 2.391648 | 1.93E-17 | *LAMA4* | 1.155406 | 2.67E-14 |
| *MMP2* | 2.37447 | 2.2E-17 | *COL6A2* | 1.598649 | 3.54E-14 |
| *DACT1* | 1.197408 | 2.22E-17 | *EFEMP1* | 1.796197 | 3.62E-14 |
| *EVC* | 1.22319 | 2.33E-17 | *AEBP1* | 1.917439 | 3.65E-14 |
| *ACTA2* | 1.914692 | 2.84E-17 | *MEDAG* | 2.042175 | 3.72E-14 |
| *GAS7* | 1.289517 | 3.47E-17 | *FILIP1L* | 1.280549 | 4.18E-14 |
| *COL3A1* | 2.520224 | 4.58E-17 | *CCN2* | 1.934882 | 4.27E-14 |
| *SERPINF1* | 2.003177 | 5E-17 | *MFAP4* | 1.886944 | 4.36E-14 |
| *GNB4* | 1.079462 | 5.25E-17 | *SSC5D* | 1.243839 | 4.46E-14 |
| *TGFB3* | 1.755902 | 5.73E-17 | *GLIPR2* | 1.245947 | 4.56E-14 |
| *GPNMB* | 1.989968 | 5.99E-17 | *C1S* | 1.747892 | 4.96E-14 |
| *THBS1* | 2.322713 | 5.99E-17 | *PRSS23* | 1.168927 | 6E-14 |
| *GLIPR1* | 1.142906 | 6.69E-17 | *NTM* | 1.345661 | 6.55E-14 |
| *ZCCHC24* | 1.405358 | 6.76E-17 | *FPR1* | 1.443118 | 6.64E-14 |
| *PDLIM3* | 1.313843 | 6.76E-17 | *SFRP4* | 2.293225 | 7.41E-14 |
| *THBS2* | 2.469537 | 6.96E-17 | *LRRC17* | 1.172025 | 7.43E-14 |

| | | | | | |
|---|---|---|---|---|---|
| *F13A1* | 1.946628 | 7.51E-17 | *LY96* | 1.306593 | 7.94E-14 |
| *CAMSAP3* | 1.211996 | 7.52E-17 | *TMEM238* | 1.568243 | 7.98E-14 |
| *IL1R1* | 1.625894 | 7.79E-17 | *C1R* | 1.514589 | 8.82E-14 |
| *EDNRA* | 1.51914 | 7.79E-17 | *CPXM2* | 1.369921 | 9.55E-14 |
| *SGCD* | 1.09362 | 7.79E-17 | *RAB3IL1* | 1.005619 | 9.69E-14 |
| *PDPN* | 1.732546 | 8.3E-17 | *KCNE4* | 1.294427 | 1.05E-13 |
| *CAVIN1* | 1.300185 | 8.3E-17 | *WIPF1* | 1.166234 | 1.09E-13 |
| *DPYSL3* | 1.573631 | 9.77E-17 | *TMEM184A* | 1.136091 | 1.12E-13 |
| *ASPN* | 2.07737 | 9.88E-17 | *COL15A1* | 1.500634 | 1.22E-13 |
| *SPOCK1* | 1.605792 | 1.17E-16 | *MYLK* | 1.324617 | 1.24E-13 |
| *ANTXR1* | 1.933463 | 1.17E-16 | *ITGA5* | 1.387628 | 1.27E-13 |
| *FMO1* | 1.176788 | 1.54E-16 | *ROR2* | 1.072725 | 1.29E-13 |
| *HEG1* | 1.290124 | 1.73E-16 | *BGN* | 1.504274 | 1.52E-13 |
| *ZNF469* | 1.071182 | 1.8E-16 | *FGF7* | 1.345607 | 1.55E-13 |
| *SVEP1* | 1.30705 | 1.82E-16 | *ELN* | 1.588643 | 1.56E-13 |

**Supplementary Table 5.** The top 200 glucose metabolism-correlated genes in the proteomics cohort.

| Gene symbol | logFC | adj.P-Val | Gene symbol | logFC | adj.P-Val |
|---|---|---|---|---|---|
| *HBA2* | 1.518319 | 1.86E-07 | *FKBP11* | 1.623657 | 5.95E-06 |
| *HBA1* | 1.575869 | 1.86E-07 | *MTARC2* | 1.594271 | 6.02E-06 |
| *P4HB* | 1.788522 | 2.26E-07 | *SLC4A4* | 2.045488 | 6.62E-06 |
| *AMY2A* | 2.046344 | 2.62E-07 | *FMNL1* | 1.736639 | 7.02E-06 |
| *AMY2B* | 2.508841 | 2.72E-07 | *NDUFA2* | 1.540236 | 7.22E-06 |
| *TUBB* | 1.927868 | 3.71E-07 | *GIPC1* | 2.218894 | 7.56E-06 |
| *CPB1* | 3.756637 | 3.71E-07 | *SLC25A22* | 1.517512 | 8.93E-06 |
| *CPA1* | 5.187461 | 3.94E-07 | *ARSL* | 1.502851 | 9.36E-06 |
| *SND1* | 1.571858 | 4.09E-07 | *MGST1* | 1.524212 | 9.55E-06 |
| *PNLIP* | 3.919663 | 4.09E-07 | *IMPA2* | 1.912337 | 9.55E-06 |
| *MUC5AC* | 1.833319 | 4.09E-07 | *NRCAM* | 1.553444 | 9.68E-06 |
| *CEL* | 3.591888 | 4.09E-07 | *CTRL* | 1.522063 | 1.01E-05 |
| *CA1* | 3.190163 | 4.09E-07 | *ELAPOR1* | 1.506479 | 1.31E-05 |
| *ALDH1A1* | 1.94254 | 4.09E-07 | *V-kappa-3* | 1.534102 | 1.34E-05 |
| *PDIA2* | 3.387334 | 4.15E-07 | *HLA-DQA1* | 1.58888 | 1.45E-05 |
| *GATM* | 3.622489 | 4.15E-07 | *CBR4* | 1.78567 | 1.63E-05 |
| *KRT6A* | 1.61595 | 4.34E-07 | *NEDD4* | 1.642896 | 1.63E-05 |
| *CTRB2* | 7.030082 | 4.52E-07 | *CLYBL* | 1.615257 | 1.83E-05 |
| *PRSS1* | 3.704339 | 4.67E-07 | *NUP155* | 1.563381 | 1.94E-05 |
| *SPTA1* | 1.569657 | 4.79E-07 | *MCFD2* | 2.611303 | 1.96E-05 |
| *TPM3* | 1.840195 | 4.87E-07 | *SLC35B2* | 1.520086 | 1.99E-05 |
| *PRSS2* | 5.638633 | 4.92E-07 | *TMEM33* | 1.527714 | 2.05E-05 |
| *IGHA1* | 2.061446 | 5.5E-07 | *CTH* | 1.791401 | 2.06E-05 |
| *IDH2* | 1.539545 | 5.5E-07 | *HSD17B8* | 1.655564 | 2.41E-05 |
| *LRRC59* | 1.512611 | 5.52E-07 | *CRABP2* | 1.503951 | 2.41E-05 |
| *HBG1* | 2.641399 | 5.62E-07 | *ACADL* | 1.647697 | 2.44E-05 |
| *ACTB* | 2.292788 | 5.62E-07 | *IER3IP1* | 1.614907 | 2.46E-05 |
| *CA2* | 1.798814 | 6.56E-07 | *EIF2B4* | 1.620973 | 2.94E-05 |
| *HBG2* | 2.793506 | 6.84E-07 | *SNAPIN* | 1.530803 | 3.04E-05 |
| *CPA2* | 2.563636 | 6.89E-07 | *PSMB5* | 1.602276 | 3.14E-05 |
| *RACK1* | 1.612703 | 6.96E-07 | *HSPA12B* | 1.951405 | 3.22E-05 |
| *NIBAN1* | 1.504244 | 7.27E-07 | *REEP6* | 2.917434 | 3.45E-05 |
| *CLIC1* | 1.500934 | 7.27E-07 | *PIK3AP1* | 1.803297 | 3.5E-05 |
| *ACAT1* | 1.65585 | 7.27E-07 | *TXNRD2* | 1.649702 | 3.71E-05 |
| *SLC4A1* | 1.673193 | 7.97E-07 | *DDT* | 1.562978 | 3.74E-05 |
| *REG1A* | 3.044204 | 7.97E-07 | *EPHX2* | 1.951049 | 3.79E-05 |
| *EPHX1* | 2.126253 | 7.97E-07 | *SGSH* | 1.597904 | 3.98E-05 |
| *AGR2* | 1.645803 | 7.97E-07 | *IgH* | 2.812334 | 4E-05 |
| *CTRC* | 3.568562 | 8.01E-07 | *CELA3A* | 1.919521 | 4.04E-05 |

| | | | | | |
|---|---|---|---|---|---|
| *PNLIPRP1* | 2.039263 | 9.71E-07 | *VPS51* | 1.51339 | 4.19E-05 |
| *GSTA2* | 2.167727 | 9.75E-07 | *SRA1* | 1.769373 | 4.19E-05 |
| *ALDH6A1* | 1.531319 | 1.12E-06 | *DEFA1* | 1.85491 | 4.27E-05 |
| *PHGDH* | 1.919513 | 1.16E-06 | *PLCG2* | 1.623802 | 4.32E-05 |
| *CELA3B* | 2.185833 | 1.23E-06 | *SYTL1* | 1.539334 | 4.61E-05 |
| *RPS7* | 2.28993 | 1.26E-06 | *ARMC8* | 1.603812 | 5.12E-05 |
| *ALDH9A1* | 1.648013 | 1.27E-06 | *RBP2* | 2.137855 | 5.26E-05 |
| *CELA2A* | 5.036042 | 1.28E-06 | *PNKP* | 1.559299 | 5.41E-05 |
| *ABAT* | 1.815832 | 1.36E-06 | *SLC39A14* | 1.629081 | 5.92E-05 |
| *AOX1* | 1.589667 | 1.36E-06 | *ECHDC3* | 1.586911 | 6.05E-05 |
| *SEC22B* | 1.776135 | 1.44E-06 | *HLA-DRB1* | 2.266455 | 6.11E-05 |
| *PTI-1* | 1.542613 | 1.46E-06 | *TRIM56* | 1.70727 | 6.34E-05 |
| *SEC61A1* | 1.984137 | 1.47E-06 | *SMPDL3B* | 1.773659 | 6.4E-05 |
| *LGALS2* | 1.901498 | 1.47E-06 | *TSTD1* | 1.635506 | 7.51E-05 |
| *PRDX2* | 2.388614 | 1.49E-06 | *RRM1* | 1.673746 | 7.82E-05 |
| *CLPS* | 3.083303 | 1.55E-06 | *PPIF* | 1.560508 | 8.14E-05 |
| *RAB3D* | 1.737744 | 1.59E-06 | *GCAT* | 2.265745 | 8.72E-05 |
| *MYO6* | 1.679121 | 1.65E-06 | *TMEM70* | 1.767277 | 8.93E-05 |
| *DLD* | 1.739716 | 1.67E-06 | *SIRT3* | 1.64034 | 8.93E-05 |
| *REG1B* | 2.456672 | 1.71E-06 | *MRPL58* | 2.312592 | 9.16E-05 |
| *CHID1* | 1.673664 | 1.73E-06 | *SYCN* | 1.850613 | 9.63E-05 |
| *ASAH1* | 1.669292 | 1.76E-06 | *NBEA* | 1.534165 | 0.000107 |
| *RPL18* | 1.568669 | 1.81E-06 | *ARSB* | 2.240146 | 0.000108 |
| *hCG_1639753* | 3.109603 | 1.85E-06 | *STEAP4* | 1.57383 | 0.000114 |
| *PSAT1* | 1.827253 | 1.86E-06 | *DPM3* | 1.863167 | 0.000136 |
| *GOT2* | 2.856613 | 1.87E-06 | *GNL3* | 3.72357 | 0.000141 |
| *STT3A* | 1.581546 | 2.01E-06 | *WDR91* | 1.501974 | 0.000148 |
| *HIBCH* | 1.502993 | 2.12E-06 | *B3GNT7* | 1.544071 | 0.000149 |
| *REG3A* | 5.376078 | 2.15E-06 | *FAM177A1* | 2.499708 | 0.00016 |
| *SERPINB3* | 1.753159 | 2.24E-06 | *LAMA1* | 2.058612 | 0.000176 |
| *GNAS* | 1.522708 | 2.26E-06 | *TP53I11* | 2.698882 | 0.00019 |
| *CRAT* | 1.79531 | 2.45E-06 | *MRPL14* | 1.629242 | 0.00019 |
| *ERLIN1* | 1.851842 | 2.48E-06 | *C1GALT1* | 1.563035 | 0.000196 |
| *FABP1* | 2.188117 | 2.55E-06 | *TMUB1* | 1.812711 | 0.000223 |
| *PBLD* | 1.510841 | 2.58E-06 | *ACAD10* | 1.790923 | 0.000227 |
| *SERPINI2* | 1.923752 | 2.6E-06 | *RASA4* | 1.579511 | 0.000234 |
| *SARDH* | 1.886059 | 2.63E-06 | *WDR5* | 1.711962 | 0.000237 |
| *CAMK2B* | 1.715353 | 2.63E-06 | *HEATR3* | 2.133554 | 0.000243 |
| *GNMT* | 1.586816 | 2.72E-06 | *USP24* | 2.673536 | 0.000265 |
| *GALE* | 1.548366 | 2.72E-06 | *PHKB* | 1.677289 | 0.000301 |
| *GP2* | 1.604078 | 2.73E-06 | *PYCR2* | 1.612255 | 0.000302 |
| *ERP27* | 2.037428 | 2.83E-06 | *CLN5* | 2.217648 | 0.000335 |
| *ERO1B* | 1.585562 | 2.85E-06 | *PAIP2B* | 1.546377 | 0.000357 |
| *PAFAH1B3* | 1.741402 | 2.92E-06 | *COX7C* | 1.818646 | 0.000371 |

| | | | | | |
|---|---|---|---|---|---|
| *HMOX2* | 2.197031 | 3.54E-06 | *ABCC4* | 1.518606 | 0.000374 |
| *MPI* | 1.525988 | 3.68E-06 | *TFF1* | 1.644083 | 0.000418 |
| *CELA2B* | 3.386187 | 3.99E-06 | *OMA1* | 1.612223 | 0.00042 |
| *ALDH1A3* | 1.616915 | 4.06E-06 | *SLC44A3* | 2.06797 | 0.000423 |
| *ITPA* | 3.94064 | 4.08E-06 | *TPST2* | 1.782206 | 0.000442 |
| *TMEM97* | 3.534161 | 4.28E-06 | *ALG8* | 1.75088 | 0.000475 |
| *EPB42* | 1.580833 | 4.29E-06 | *RHAG* | 2.185727 | 0.000504 |
| *hCG_18987* | 1.771572 | 4.31E-06 | *STRIP1* | 1.576428 | 0.000561 |
| *KLK1* | 2.139192 | 4.47E-06 | *KCTD14* | 2.295296 | 0.00057 |
| *KPNA3* | 1.879586 | 4.5E-06 | *OSTC* | 1.666058 | 0.000606 |
| *VPS25* | 1.574701 | 4.63E-06 | *IDNK* | 1.680605 | 0.000624 |
| *GAMT* | 1.643968 | 4.83E-06 | *FLYWCH2* | 1.737089 | 0.000658 |
| *CBX5* | 1.640728 | 4.86E-06 | *SNX17* | 1.963776 | 0.000666 |
| *ALDOB* | 1.595867 | 5.03E-06 | *PRG4* | 2.315483 | 0.000833 |
| *PLA2G1B* | 7.414452 | 5.45E-06 | *ATF6B* | 2.315584 | 0.000938 |
| *SGPL1* | 2.220681 | 5.48E-06 | *IF1IC2* | 1.646366 | 0.000984 |
| *SGCD* | 2.077625 | 5.95E-06 | *BTAF1* | 1.86243 | 0.001131 |

**Supplementary Figures**

**Supplementary Figure 1.** Functional and pathway annotation of tumor matrix-correlated genes and glucose metabolism-correlated genes. (A) The top 30 KEGG analysis terms from tumor matrix-correlated genes. (B) The top 20 most enriched KEGG pathways in glucose metabolism-correlated genes.

**Supplementary Figure 2.** Expression and clinical correlations of CLIC1 in PDAC tissues. (A) Univariate Cox regression analysis of clinicopathological parameters for overall survival based on the TMA cohort. (B) Respective sample scoring on the staining of CLIC1 expression in KPC mice.

**Supplementary Figure 3.** Expression and growth-promoting ability of CLIC1 in PDAC. (A-B) Expression of CLIC1 at mRNA and protein levels in various PDAC cell lines. (C-D) Efficiency of interference at mRNA and protein levels of CLIC1 expression in PANC-1 and PATU8988 cells treated with shRNA (mean ± SEM., two-tailed unpaired *t* test). (E-F) Relative mRNA and protein expression of CLIC1 in CLIC1$^{OE}$ cells (Capan-1 and SW-1990) (mean ± SEM., two-tailed unpaired *t* test). (G-J) Immunohistochemical staining images of PCNA and Ki67 from the tumor tissues in subcutaneous and orthotopic xenograft models. Scale bar: 50 μm. ***P* < 0.01, ****P* < 0.001.

**Supplementary Figure 4.** CLIC1 promotes the growth of PDAC cells by enhancing the Warburg effect. (A) The correlation between CLIC1 expression and the HALLMARK_GLYCOLYSIS gene set in PDAC samples analyzed by GSEA based on data from the GEO and TCGA databases. NES, normalized enrichment score. (B) The correlation between *CLIC1* expression and two commonly used glycolysis-related gene sets (HALLMARK_GLYCOLYSIS and KEGG_GLYCOLYSIS_GLUCONEOGENESIS) from TCGA databases. (C-D) Relative mRNA levels of glycolysis-relevant genes in CLCI1$^{OE}$ and CLCI1$^{KD}$ cells (mean ± SEM., two-tailed unpaired *t* test). (E) Evaluation of the glycolysis rate of CLCI1$^{KD}$ and CLCI1$^{OE}$ cells as measured by ECAR. (F-G) Relative glucose uptake and lactate production ability in the CLIC1 knockdown and overexpression groups and the corresponding control groups (mean ± SEM., two-tailed unpaired *t* test). (H-I) Representative IHC and corresponding CT and PET-CT images of the CLIC1 low- and high- expression groups and the comparison of SUV-Max between the two groups (n = 7 cases per group, mean ± SEM., two-tailed unpaired *t* test, scale bar: 50 μm). (J-K) Colony formation ability of CLCI1$^{OE}$ cells treated with 2-DG and galactose (mean ± SEM., one-way ANOVA with Tukey's multiple comparison test). *$P < 0.05$, **$P < 0.01$, ***$P < 0.001$.

Supplementary Figure 1

A

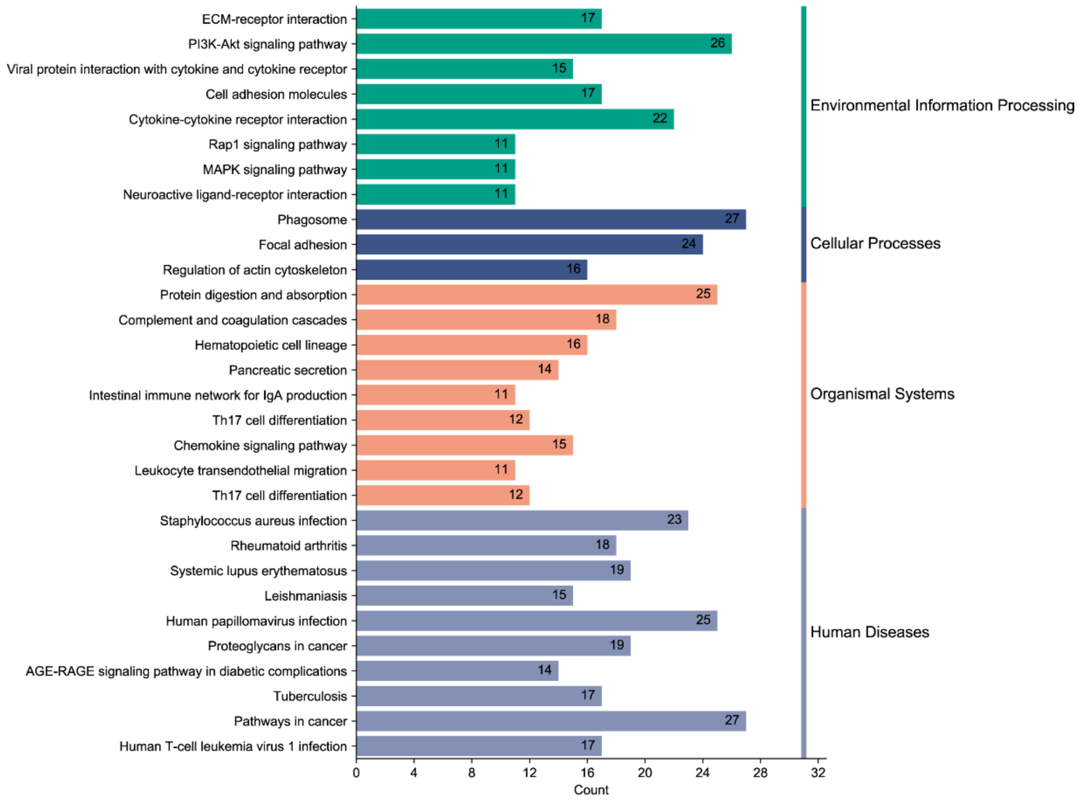


B

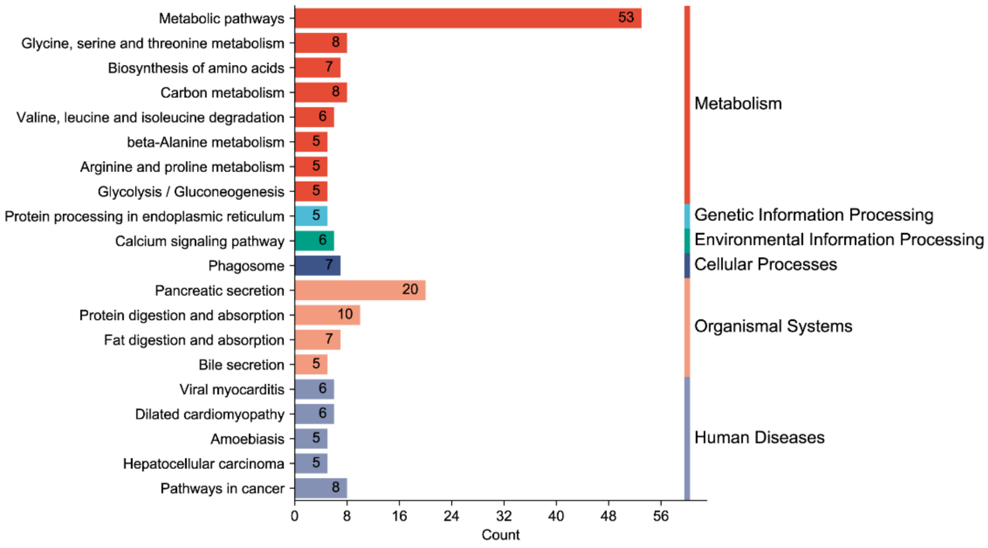

Supplementary Figure 2

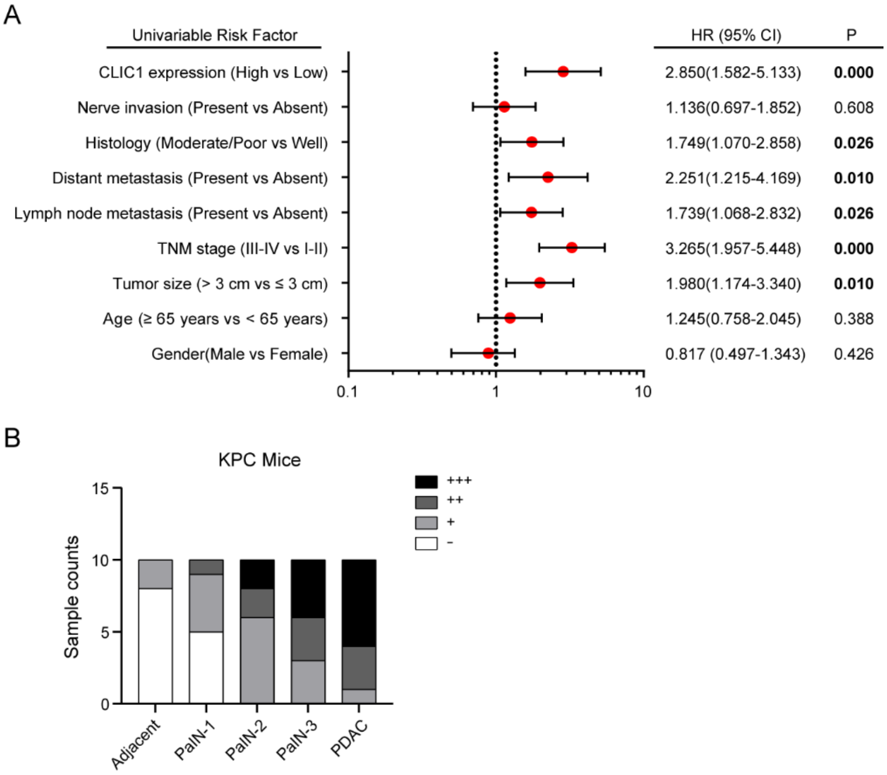

Supplementary Figure 3

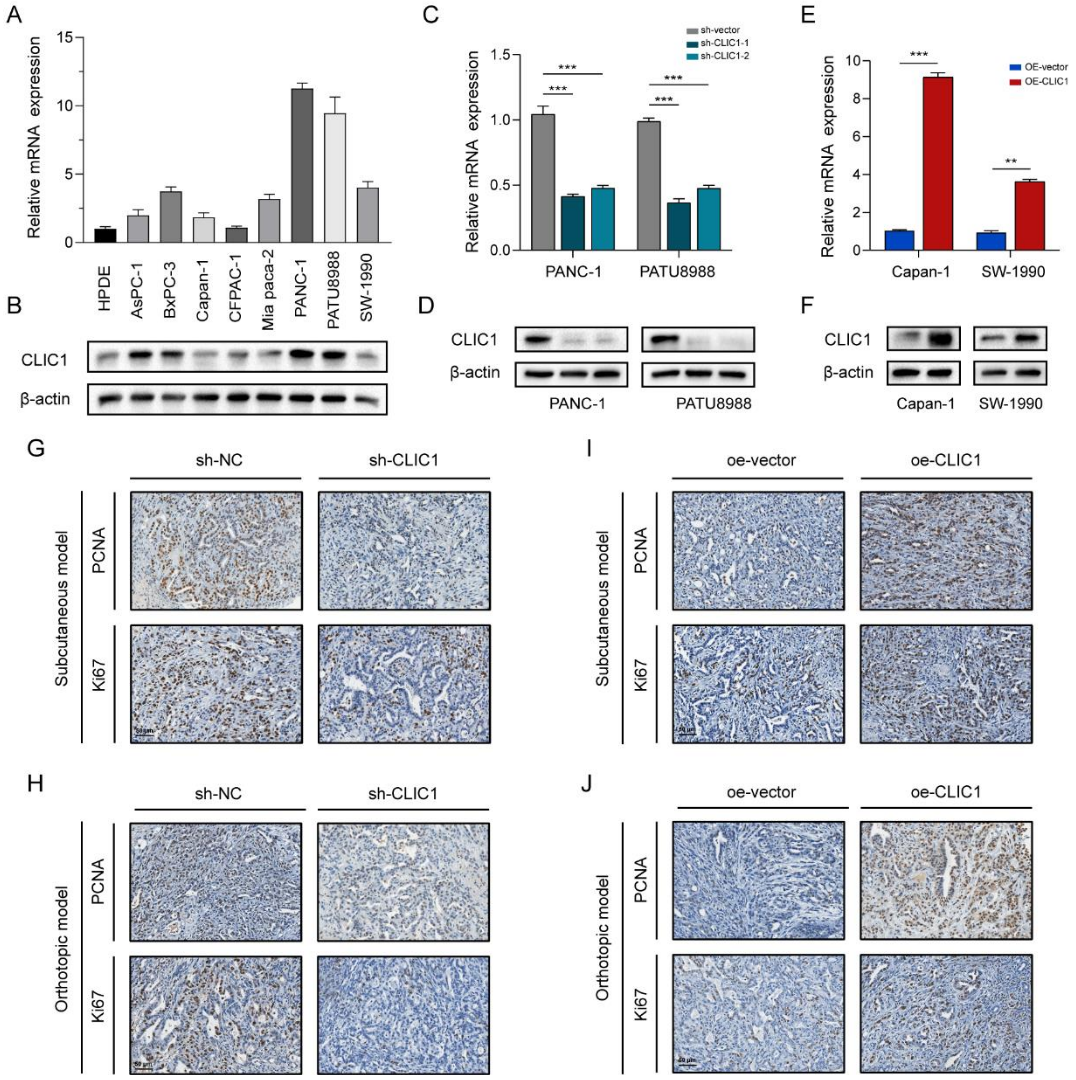

Supplementary Figure 4

A

| Datasets | NES | FDR q-value | Nom p-value |
|---|---|---|---|
| TCGA | 2.3348 | 0.0000 | 0.0000 |
| GSE 15471 | 2.4317 | 0.0000 | 0.0000 |
| GSE 28735 | 2.3328 | 0.0000 | 0.0000 |
| GSE 16515 | 2.5203 | 0.0000 | 0.0000 |
| GSE 102238 | 2.1003 | 0.0000 | 0.0000 |

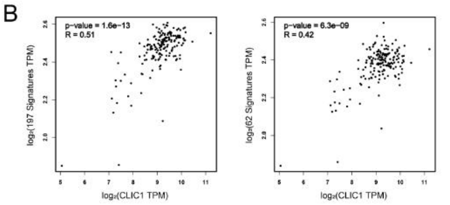


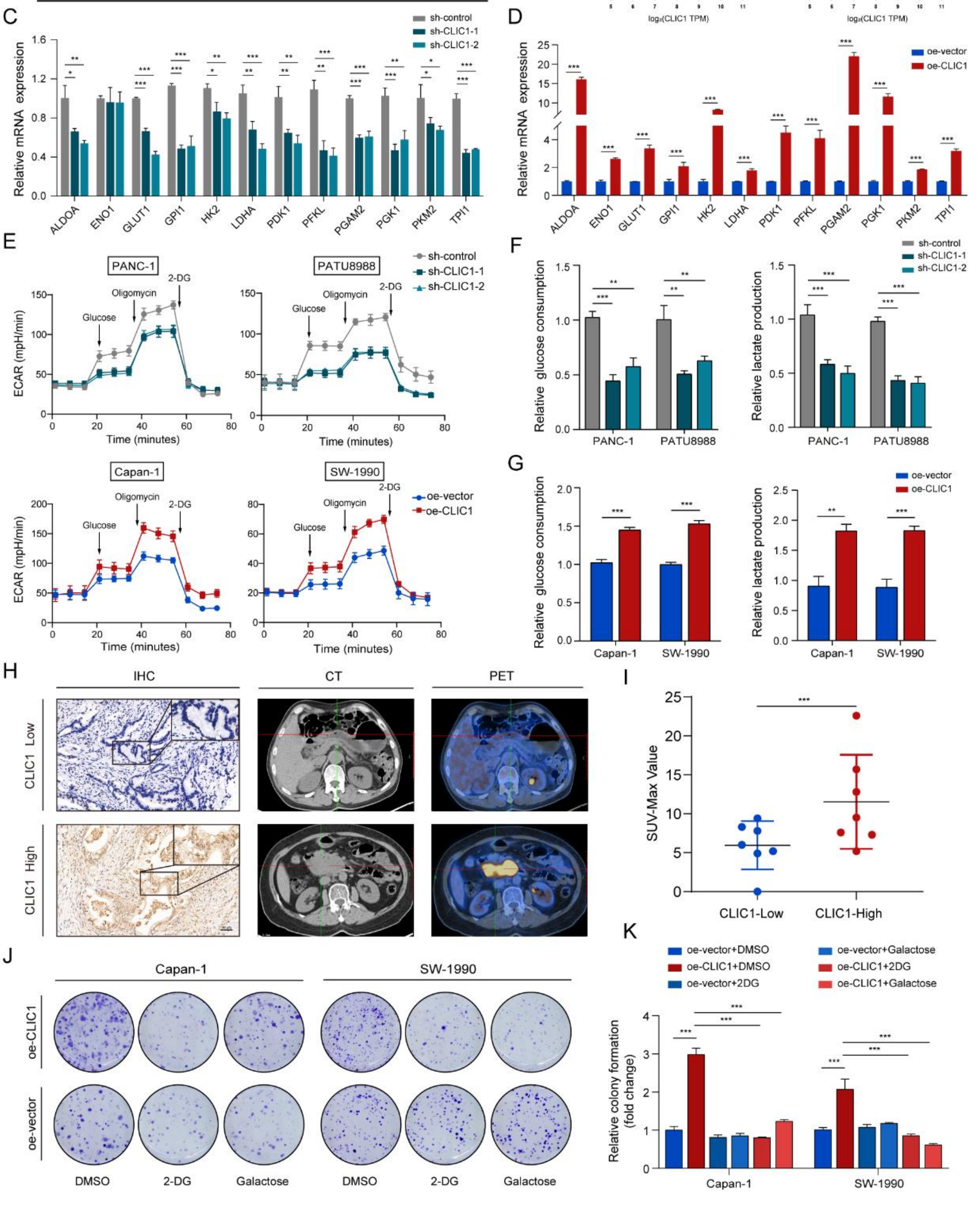
